# Stem cell derived astrocytes with *POLG* mutations and mitochondrial dysfunction including abnormal NAD+ metabolism is toxic for neurons

**DOI:** 10.1101/2020.12.20.423652

**Authors:** Kristina Xiao Liang, Atefeh Kianian, Anbin Chen, Cecilie Katrin Kristiansen, Yu Hong, Jessica Furriol, Lena Elise Høyland, Mathias Ziegler, Torbjørn Kråkenes, Charalampos Tzoulis, Gareth John Sullivan, Laurence A. Bindoff

**Author notes:** These first authors contributed equally (Kristina Xiao Liang, Atefeh Kianian). These authors share corresponding authorship (Kristina Xiao Liang, Gareth John Sullivan, Laurence A. Bindoff). Corresponding author, Kristina Xiao Liang, Neuro-SysMed, Department of Neurology, Haukeland University Hospital, PO Box 5021, Bergen, Norway., Co-corresponding author, Laurence A. Bindoff, Neuro-SysMed, Department of Neurology, Haukeland University Hospital, PO Box 5021, Bergen, Norway., Co-corresponding author, Gareth John Sullivan^f^, Department of Pediatric Research, Oslo University Hospital, P. O. Box 4950, Nydalen, 0424 Oslo, Norway.

## Abstract

The inability to reliably replicate mitochondrial DNA (mtDNA) by mitochondrial DNA polymerase gamma (POLG) leads to a subset of common mitochondrial diseases associated with neuronal death and depletion of neuronal mtDNA. Defining disease mechanisms remains difficult due to the limited access to human tissue. Astrocytes are highly abundant in the brain, playing a crucial role in the support and modulation of neuronal function. Astrocytes also respond to insults affecting the brain. Following damage to the center neural system, which can be hypoxia, inflammation or neurodegeneration, astrocytes become activated and lose their supportive role and gain toxic functions that induce rapid death of neurons and oligodendrocytes. The role of astrocyte reactivation and the consequences this has for neuronal homeostasis in mitochondrial diseases has not been explored. Here, using patient cells carrying *POLG* mutations, we generated iPSCs and then differentiated into astrocytes. We demonstrated that POLG-astrocytes exhibited both mitochondrial dysfunctions, including loss of mitochondrial membrane potential, energy failure, complex I and IV defects, disturbed NAD^+^/NADH metabolism, and mtDNA depletion. Further, POLG derived astrocytes presented an A1-like reactive phenotype with increased proliferation, invasion, upregulation of pathways involved in response to stimulus, immune system process, cell proliferation and cell killing. Under direct and indirect co-culture with neurons, POLG-astrocytes exhibited a toxic effect leading to the death of neurons. Our findings demonstrate that mitochondrial dysfunction caused by *POLG* mutations leads not only to intrinsic defects in energy metabolism affecting both neurons and astrocytes, but also to neurotoxic damage driven by astrocytes. Our studies provide a robust astroglia-neuronal interaction model for future investigation of mitochondrial involvement in neurogenesis and neurodegenerative diseases.

**Highlights:**

▪ Patient-specific astrocytes harbouring a *POLG* mutation showed lower mitochondrial membrane potential and mtDNA depletion.
▪ POLG-astrocytes generated elevated L-lactate as the end glycolytic product.
▪ Patient-specific astrocytes with *POLG* mutations exhibited mitochondrial respiratory chain disruption accompanied with abnormal UCP2/SirT1/SirT3 mediated NAD^+^ metabolism.
▪ Suppressed complex I and IV-driven respiration contributed to the pathological mechanisms in POLG-related disease.
▪ POLG-astrocytes exhibit A1-reactive phenotype and neurotoxic potential.

**Graphic abstract:** 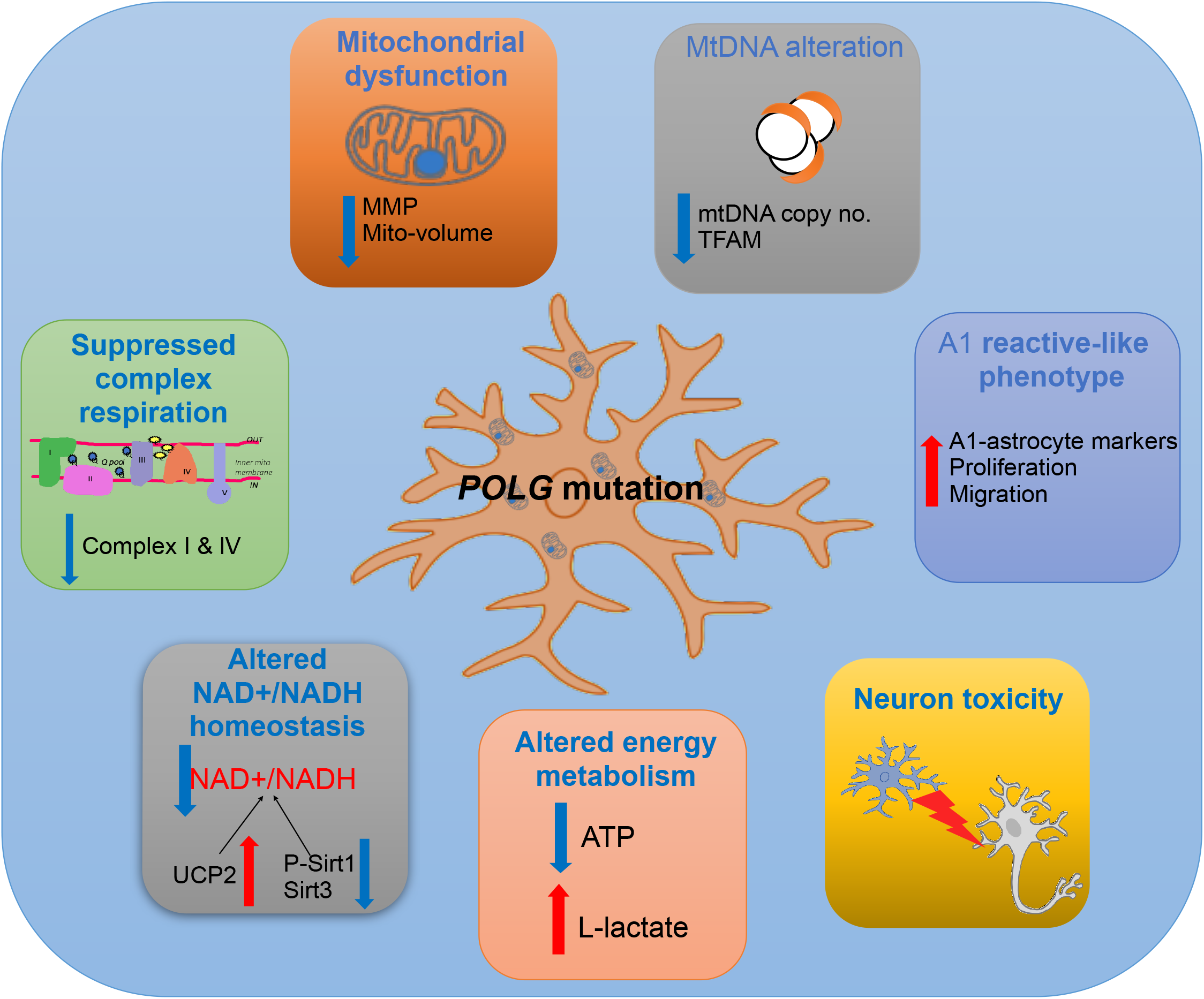

## Summary

The *POLG* gene encodes the catalytic subunit of polymerase gamma (pol γ), the enzyme that replicates and repairs mitochondrial DNA (mtDNA) [1]. Mutations in *POLG* are the most common cause of inherited mitochondrial disease and clinically these demonstrate a continuum of overlapping phenotypes from infantile disorders. At the molecular level, mutations in *POLG* lead to mtDNA maintenance defects and mitochondrial dysfunction. Interestingly, the molecular mtDNA defects differ depending on the tissue; multiple deletions are seen in skeletal muscle while neurons and hepatocytes show mtDNA depletion [2]. The mechanisms behind decreased mtDNA quality and mitochondrial dysfunction in POLG related disease remain, however, elusive. In post-mortem studies, we have shown that the loss of neurons in POLG disease is driven by severe mtDNA depletion [2]. How this influences mitochondrial function, including energy metabolism and redox state, and how this causes cell death, remain areas of intense speculation.

The contribution of glial cell dysfunction to disease has come more into focus [3–5]. Astrocytes are highly abundant [6] and play a crucial role in the support and modulation of neuronal function including regulating glutamate turnover, ion and water homeostasis, synapse formation and modulation, tissue repair, energy storage, and defense against oxidative stress [6,7]. These cells are also critical for neuronal metabolism [8,9] and are a major component of neurovascular coupling [10]. Greater understanding of the importance of astrocytes has shifted focus to the role of these cells in neurological diseases such as Parkinson’s disease (PD) [11], Alzheimer’s disease (AD) [12,13], Huntington’s disease (HD) [14] and amyotrophic lateral sclerosis (ALS) [15]. However, the role of astrocytes in mitochondrial diseases such as POLG has yet to be explored.

Both cerebral injury and disease can induce astrocytes to enter a ‘reactive’ state in which gene expression changes markedly [5].These reactive astrocytes are classified into A1 and A2 types according to their function: A1 astrocytes always lose their supportive role and gain toxic functions modulated via secreted neurotoxins that induce rapid death of neurons and oligodendrocytes; A2 astrocytes promote neuronal survival and tissue repair. Recent studies suggested that activated microglia induced astrocytes conversion to the A1 reactive phenotype by releasing Interleukin 1 alpha (IL1α), Tumor Necrosis Factor alpha (TNFα), and the Complement Component Subunit 1q (C1q) [5]. In neurodegenerative states such as AD [16], PD [17], multiple sclerosis (MS) [16] and ALS [18], reactive astrocytes can display both neuroprotective and neurodegenerative functions. The role of astrocyte reactivation and the consequences this has for neuronal homeostasis in mitochondrial diseases has not been explored.

The aim of this study was to examine how glial cells contribute to the pathophysiology of POLG-related disease. To do this, we generated patient-specific iPSC-derived astrocytes from two patients, one homozygous for c.2243G>C; p.W748S (WS5A) and one compound heterozygous c.1399G>A/c.2243G>C; p.A467T/W748S (CP2A). We found that POLG-astrocytes displayed multiple disease phenotypes including the loss of mitochondrial volume and membrane potential, loss of both complex I and IV, mtDNA depletion, ATP depletion and elevated L-lactate production. Further, we found defective NAD metabolism with downregulation of phosphorylation of Sirtuin 1 and Sirtuin 3 and upregulation of UCP2. POLG-astrocytes also displayed a reactivate-like phenotype with increased cell proliferation, migration, and upregulated A1 reactive astrocyte related gene expression. Moreover, in co-culture experiments, we observed a significant reduction in the number of normal neurons in the presence of POLG-astrocytes, as well as the accumulation of apoptotic cells. Overall, our findings indicate that dysfunctional astrocytes contribute to the pathogenesis of POLG disease. This will have major implications for any future intervention and treatment of POLG-related disorders.

## Results

### Generation and characterization of control and patient-specific astrocytes carrying *POLG* mutations

We generated patient iPSCs (Table S1) as previously described [19] using fibroblasts of two patients, one homozygous for c.2243G>C; p.W748S (WS5A) and one compound heterozygous c.1399G>A/c.2243G>C; p.A467T/W748S (CP2A). Control iPSCs were generated from two normal human fibroblast lines, Detroit 551 and AG05836B. In addition, we employed two human embryonic stem cell (hESC) lines (HS429 and HS360) as controls. To minimize intra-clonal variation, multiple clones for each iPSC line were used, including 4 individual clones for Detroit 551 control line, one clone for AG05836 control and 3 independent clones each for WS5A and CP2A patient lines. A commercial human primary astrocyte line, normal human astrocytes (NHA), was used as control for the astrocyte lineage.

To generate patient and control astrocytes (Fig. 1a), we first generated neural stem cells (NSCs) using a previously published protocol [19]. Within 5 days of induction, iPSC cells (Fig. 1b, (a)) with POU5F1 expression (Fig. 1b, (e)) underwent morphological changes from a colony cell shape to become neuroepithelium with a sphere-like structure (Fig. 1b, (b)) that expressed PAX6 (Fig. 1b, (f)). Neuroepithelium was then lifted into suspension culture to generate neurospheres (Fig. 1b, (c)) with expression SOX2 (Fig. 1b, (g)) and then dissociated for further growth into NSC monolayers (Fig. 1b, (d)) with NESTIN expression (Fig. 1b, (h)). NSCs expressed the markers SOX2 and NESTIN as evaluated by immunofluorescent staining (Fig. S 1) and showed high purity with over 99% of the cells co-expressing NESTIN and PAX6 assessed by flow cytometry (Fig. 1c). Next the iPSC-derived NSCs were differentiated to astrocytes (Fig. 1d, Table S2) using a previously published protocol with some modifications [20]. The resulting astrocytes had the typical ramified shape and a stellate morphology (Fig. 1d). After 4 weeks’ differentiation, the cells were further matured in maturation medium (Table S3) for up to 3 months. We succeeded in generating astrocytes with appropriate morphology from all the described lines (Fig. S 2).

**Figure 1.**
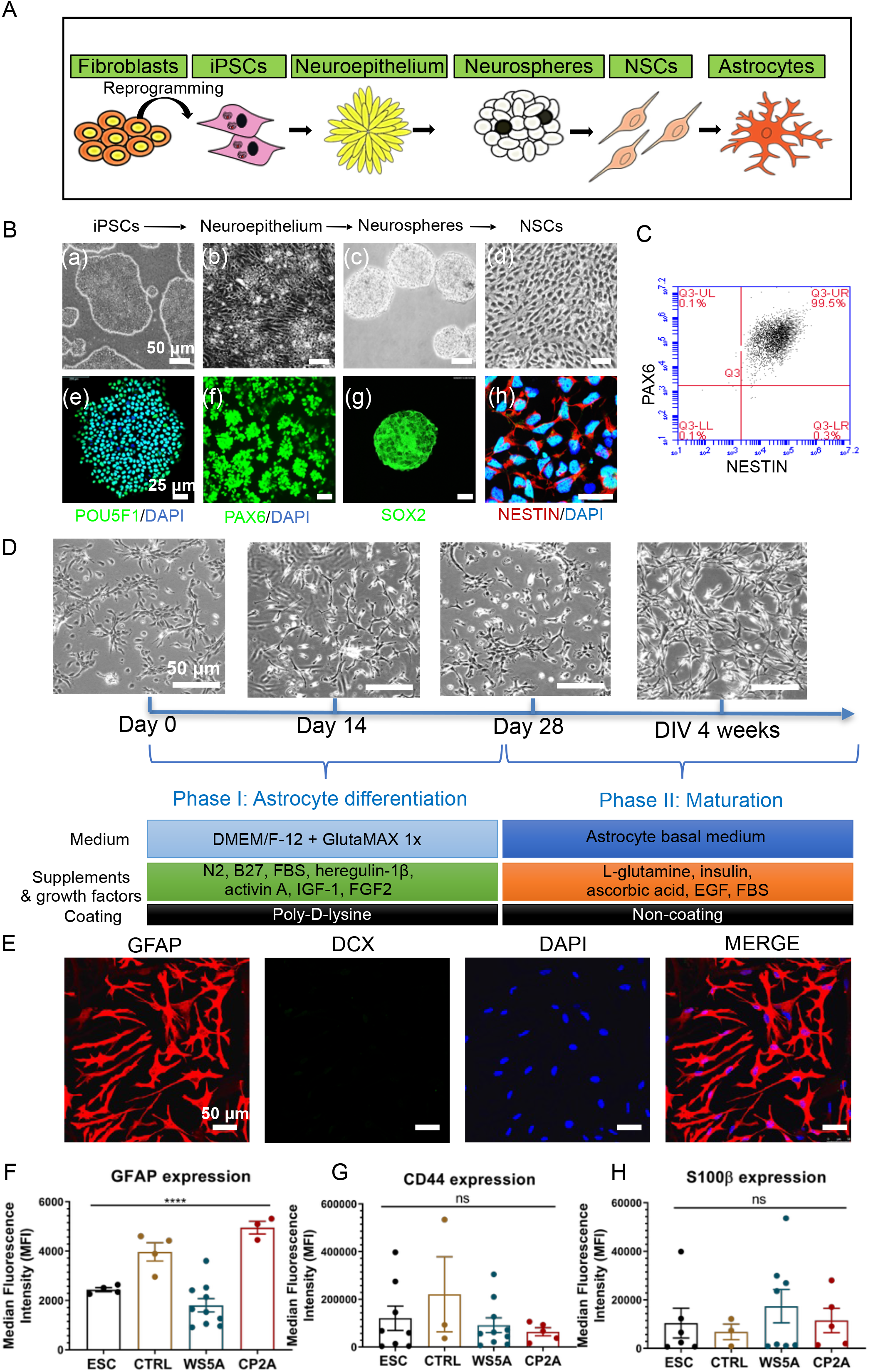
Generation and characterization of astrocytes derived from ESCs, control and POLG iPSCs. a. Flow chart of reprogramming of iPSCs and astrocytic differentiation via defined stages. b. Representative phase contrast images (upper panel) and immunostaining for specific stages during neural induction from iPSCs to NSCs. Nuclei are stained with DAPI (blue). Scale bar is 25 μm. c. Plot for the percentage of the positive staining with neural progenitor markers PAX6 and NESTIN in iPSC-derived NSCs. d. Representative phase-contrast images during 4 weeks of astrocyte differentiation from NSCs and cell culture medium information for both differentiation and maturation procedures. Scale bar is 50 μm. e. Representative phase-contrast and confocal images of immunostaining for GFAP and DCX in NHA, ESC-derived astrocytes and iPSC-derived astrocytes from control and patients. Nuclei are stained with DAPI (blue). Scale bar is 50 μm. f - h. Flow cytometric analysis for protein expression of GFAP and CD44 and S100β generated from HNA and ESC/iPSC-derived astrocytes.

We then validated the derived astrocytes using immunocytochemical and immunofluorescent staining (ICC/IF) against key astrocyte markers. We confirmed robust expression of markers for mature astrocytes including glial fibrillary acidic protein (GFAP) (Fig. 1e, Fig. S 3) and S100 calcium-binding protein β (S100β) (Fig. S 4). We detected positive DoubleCortin (DCX) expressing cells in NHA, indicating the neuronal contamination (Fig. S 3) We found no evidence of neuronal contamination in our ESC/iPSC-derived astrocytes (Fig. 1e, Fig. S 3). Astrocytic identity was further confirmed, and purity assessed by flow cytometric analysis against GFAP, CD44 and S100β antibodies. All control and patient lines exhibited 89.4%-99.9% GFAP positive cells (Fig. S 5), 89.9%-99.8% CD44 positive cells (Fig. S 6) and 80.6%-99.3% S100β positive cells (Fig. S 7). While the expression level of GFAP (Fig. 1f), CD44 (Fig. 1g) and S100β (Fig. 1h) varied in different clones, no significant differences were detected in CD44 and S100β and a significant difference was detected in GFAP expression. To examine the functional identity of the astrocytes, we assessed expression of the excitatory amino acid transporter 1 (EAAT1, also known as GLAST-1) and glutamine synthetase (GluSyn). Immunostaining demonstrated positive expression of EAAT1 and GluSyn in both NHAs and ESC/iPSC-derived astrocytes (Fig. 2a). Over 90% of the cells showed positive staining for both EAAT1 (Fig. 2b, Fig. S 8) and GluSyn (Fig. 2c, Fig. S 9) based on flow cytometric analysis. We found a different level of EAAT1 expression among different astrocytes (Fig. 2d), but similar expression of GluSyn level (Fig. 2e). We also confirmed that patient derived astrocytes carried the respective *POLG* mutations (Fig. f). These data demonstrate that both patient and control lines can generate robust populations of functional astrocyte cells. The above iPSC-derived astrocytes were used for all subsequent experiments to explore disease phenotype and mechanism.

**Figure 2.**
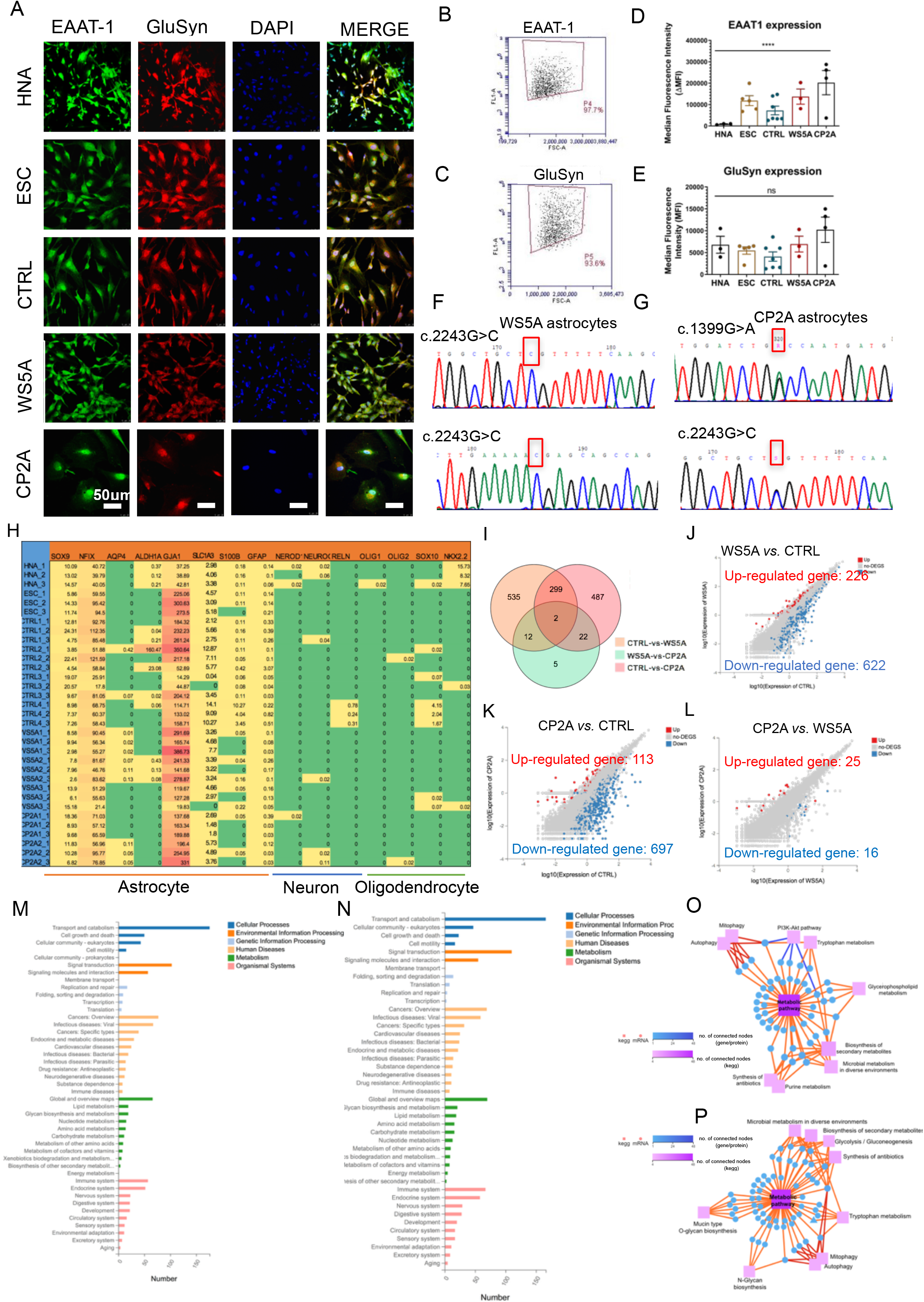
Characterization of functional markers in astrocytes derived from ESCs, control and POLG iPSCs. a. Representative confocal images of immunostaining of NHA and both control and patient astrocytes for EAAT1 (green) and GluSyn (red). Nuclei are stained with DAPI (blue). Scale bar is 50 μm. b & c. Representative plots of flow cytometric quantification by calculating the percentage of positively stained cells EAAT1 (b) and GluSyn (c). d & e. Flow cytometric measurements for protein expression of EAAT1 and GluSyn in HNA and ESC/iPSC-derived astrocytes. f. Sequencing chromatogram in WS5A and CP2A iPSC-derived astrocytes.

### Transcriptomics confirms lineage identity and metabolic effect of *POLG* mutations

We performed RNA sequencing (RNA-seq) to investigate lineage identity and potential disease-specific changes in gene expression. We compared the transcriptomes of astrocytes derived from three WS5A clones, two CP2A clones and a panel of control lines including NHAs, 2 ESCs, Detroit 551 (4 clones) and AG05836B (1 clone). Lineage identity studies showed significant homogeneity with clustering of astrocytic markers *SOX9, NFIX, AQP4, ALDH1A1, GJA1, SLC1A3, S100B* and *GFAP* in all astrocyte samples. Further, we found no expression of neuronal markers *NEUROD1, NEUROG2*, and *RELN* or oligodendrocyte markers *OLIG1, OLIG2, SOX10* and *NKX2.2* (Fig. S10). Additionally, the level of maturation of iPSC and ESC derived astrocytes was similar to primary astrocytes (Fig. S10).

Next, we investigated if there were any differentially expressed genes (DEGs) between the patient and control lines. We identified 487 significant DEGs in WS5A and 535 DEGs in CP2A derived astrocytes versus control (statistical significance cutoff was p = 0.05) (Fig. S11). Only 5 DEGs were found when comparing CP2A versus control and WS5A versus control astrocyte transcriptomes (Fig. S11). In the WS5A astrocytes, 226 genes were up-regulated while 622 genes were down-regulated compared to controls (Fig. S12) while in CP2A astrocytes, 113 genes were up-regulated and 697 genes down-regulated (Fig. S12). When comparing CP2A and WS5A astrocyte lines, only 25 genes were up-regulated while 16 genes were down-regulated (Fig. S12). KEGG pathway classifications revealed enrichment for DEGs involved in different pathways including cellular processes, environmental information processing, genetic information process, human disease, metabolism and organismal system in both patient astrocytes lines as compared to controls (Fig. S13 and S14). The DEGs in metabolism pathways included genes related to lipid metabolism, glycan biosynthesis and metabolism, nucleotide metabolism, amino acid metabolism, carbohydrate metabolism, metabolism of other amino acids and cofactors and vitamins, xenobiotics biodegradation and metabolism, biosynthesis of the secondary metabolite and energy metabolism in both patient astrocytes versus controls (Fig.S13 and S14). When comparing both patient astrocytes to controls, the following ten metabolic pathway classifications were identified, including (1) Lipid Metabolism, (2) Glycan Biosynthesis and Metabolism, (3) Nucleotide Metabolism, (4) Amino Acid Metabolism, (5) Carbohydrate Metabolism, (6) Metabolism of Other Amino Acids, (7) Metabolism of Cofactors and Vitamins, (8) Xenobiotics Biodegradation and Metabolism, (9) Biosynthesis of the Secondary Metabolite and (10) Energy Metabolism. However, only Glycan Biosynthesis and Metabolism was enriched when comparing the two patient lines (Fig. S15).

Pathway analysis of down-regulated DEGs indicated that DEGs were mainly involved in metabolic pathways in both patients compared to controls (Fig. S16). KEGG pathway enrichment analysis of down-regulated DEGs revealed enrichment for genes involved in metabolic pathways in both patient astrocytes lines as compared to controls (Fig. S17). The top ten DEGs enriched in KEGG metabolic pathways were *XYLT1, PTGS1, MGLL, ST6GALNAC5, NPR, HKDC1, CSGAL, NACT1, GALNT15, ALDH3A1* and *LTC4S* in WS5A astrocytes versus control (Table S5). In CP2A astrocytes, the top ten metabolic pathway DEGs were *ALDH1A1, PLEKHA6, B4GALNT3, ASS1, ZBTB7C, GUCY1A1, BMPER, PDE4B, GCNT3* and *PTGDS* (Table S6). These results confirm that our protocol produces astrocytes at high efficiency, which these are similar to primary human astrocytes in maturity and those astrocytes containing *POLG* mutations exhibit metabolic changes.

### POLG-astrocytes manifest mitochondrial dysfunction and mtDNA depletion

Next, we assessed mitochondrial function in patient and control astrocytes. To visualize mitochondria morphology, we used mitochondrial outer membrane import receptor subunit TOM20 (TOMM20) staining and confocal microscopy. This showed a similar mitochondrial morphology in both patient and control astrocytes (Fig. 3a). We then assessed mitochondrial volume and mitochondrial membrane potential (MMP) using double staining with MitoTracker Green (MTG) and Tetramethylrhodamine ethyl ester (TMRE) [21,22]. Both total MMP, measured using TMRE alone (Fig. 3b) and specific MMP (Fig. 3c), calculated by the ratio of TMRE/MTG were significantly lower in patient astrocytes compared to control. Mitochondrial volume, measured by MTG (Fig. 3d), showed significantly lower levels in the patient astrocytes compared to control cells. However, similar mitochondrial volume was also seen when we assessed the level of TOMM20 by flow cytometry (Fig. 3e), although the change did not reach significance. These findings suggest that *POLG* mutations lead to mitochondrial dysfunction with decreased mitochondrial volume and mitochondrial depolarization in astrocytes, without affecting mitochondrial morphology. We then assessed ATP production and L-lactate levels. Using high-performance Liquid Chromatography Mass Spectrometry (LC-MS)-based metabolomic analysis, we detected a significant drop in ATP production in both WS5A and CP2A astrocytes compared to the controls (Fig. 3f). Colorimetric analysis of L-lactate production revealed a significant elevation in patient astrocytes compared to controls (Fig. 3g).

**Figure 3.**
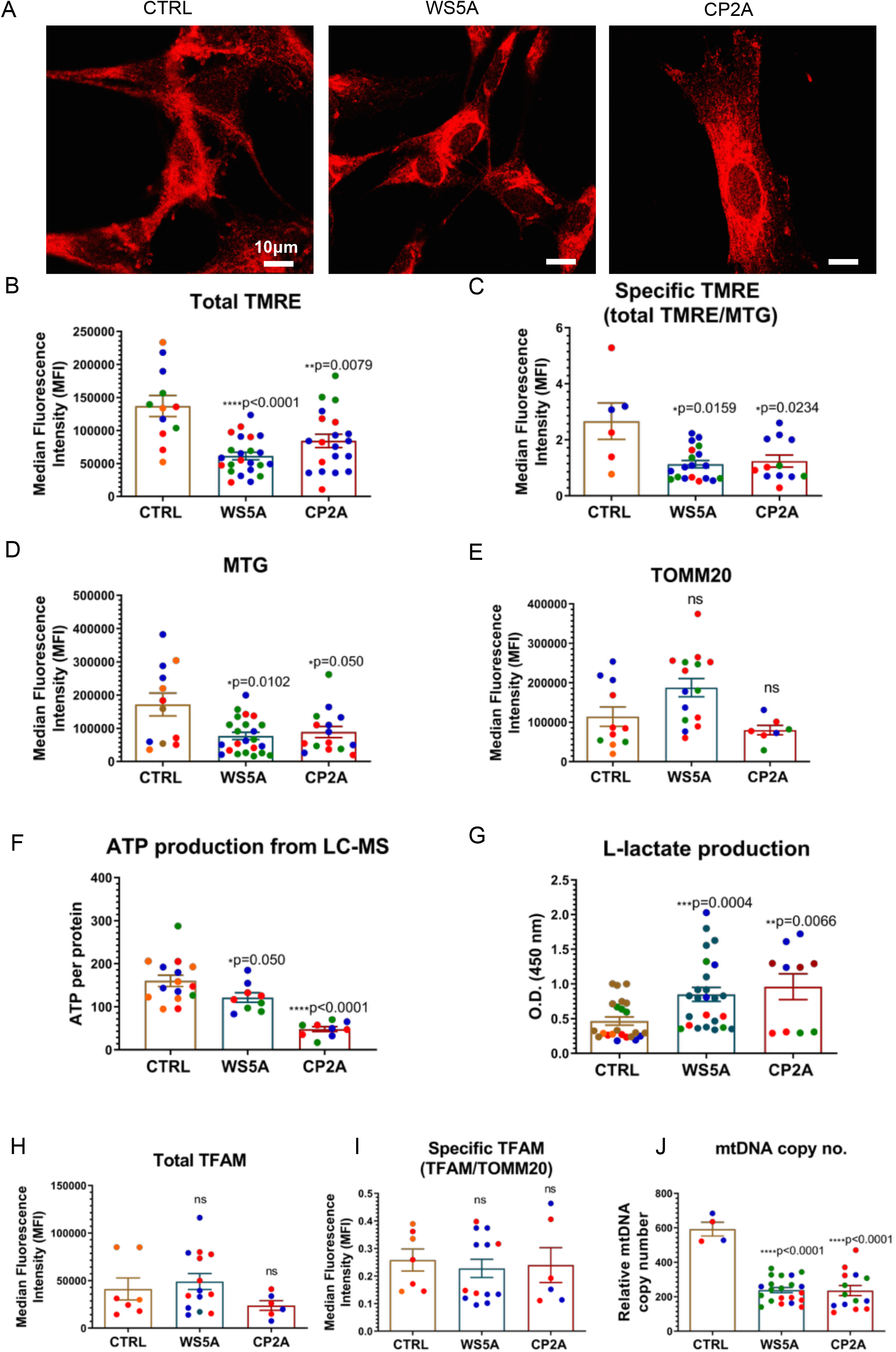
POLG-astrocytes exhibit impaired mitochondrial function, switched glycolysis and mtDNA alteration. a. Representative confocal images of control and patient astrocytes stained for TOMM20. Scale bar is 10 μm. b & c. Flow cytometric analysis for MMP at total level measured by TMRE (b) and specific MMP level calculated by total TMRE/MTG (c). d & e. Mitochondrial volume measured by MTG (d) and TOMM20 (e). f. Measurement of ATP production using LC-MS analysis. g. Colorimetric analysis for L-lactate generation. h & i: Flow cytometric analysis of TFAM protein expression at total level (h) and specific TFAM level calculated by total TFAM/TOMM20 (i). j. Relative mtDNA copy number analyzed by QPCR using primers and fluorogenic probes for regions of ND1 and nuclear gene APP.

Next, we investigated alterations in mtDNA copy number using two approaches, one an indirect method based on flow cytometry and the other direct measurement using RT-qPCR. As the mitochondrial transcription factor A (TFAM) binds mtDNA in molar quantities, we used flow cytometry to determine the level of TFAM and thus indirectly measure mtDNA content. By assessing TFAM, we detected a similar level in total TFAM protein expression (Fig. 3h) and specific level (Fig. 3i) following correction for mitochondrial volume (ratio to TOMM20) in patients versus controls. We then quantified the relative mtDNA copy number using qPCR and observed a significant mtDNA depletion in POLG patient astrocytes compared to controls (Fig. 3j). Taken together, these findings confirm that *POLG* mutations have a major impact on iPSC-derived astrocyte mitochondria with decreased mitochondrial volume, lower membrane potential and ATP production, and mtDNA depletion. These changes appear to drive a switch to greater glucose metabolism.

### POLG-astrocytes display a loss of complex I and IV

In micro-dissected post-mortem neurons, we showed previously that respiratory chain complex I was lost in POLG affected frontal and cerebellar neurons [2] and we subsequently confirmed this in iPSC-derived neural stem cells (NSCs) and iPSC-derived dopaminergic (DA) neurons [19]. We, therefore, investigated the levels of each respiratory chain complex in our astrocytes using flow cytometry, immunohistochemistry, and western blotting.

In agreement with our previous observations, we found a clear loss of complex I in patient derived astrocytes using immunostaining against NDUFB10 (Fig. 4a). Since the resolution of confocal microscopy is limited, we performed flow cytometry and western blotting to confirm our observation. Flow quantification of complex I, II and IV in astrocytes showed significantly decreased expression of complex I subunit NDFUB10 (Fig. 4b and c) and complex IV subunit 4, COX IV (Fig. 4d & e) in patient astrocytes. In contrast, complex II (Fig. 4f & g), which is wholly encoded by nuclear DNA, showed a significantly higher expression level via assessment of the SDHA subunit. Western analysis further confirmed lower complex I and IV in patient derived astrocytes (Fig. 4h & i), although WS5A astrocytes did not reach significance. These data indicate that the respiratory chain defect in astrocytes carrying *POLG* mutations appears more extensive than that seen in post-mortem neurons [23] and neuronal stem cells derived from the same iPSCs [19], where only loss of complex I was identified. A decreased expression of voltage-dependent anion channels (VDAC), located in the mitochondrial outer membrane, was also identified in WS5A and CP2A astrocytes compared to controls, although it failed to reach significance (Fig. 4h & i). Our data suggests that *POLG* mutations in astrocytes lead to loss of both mitochondrial complex I and IV.

**Figure 4.**
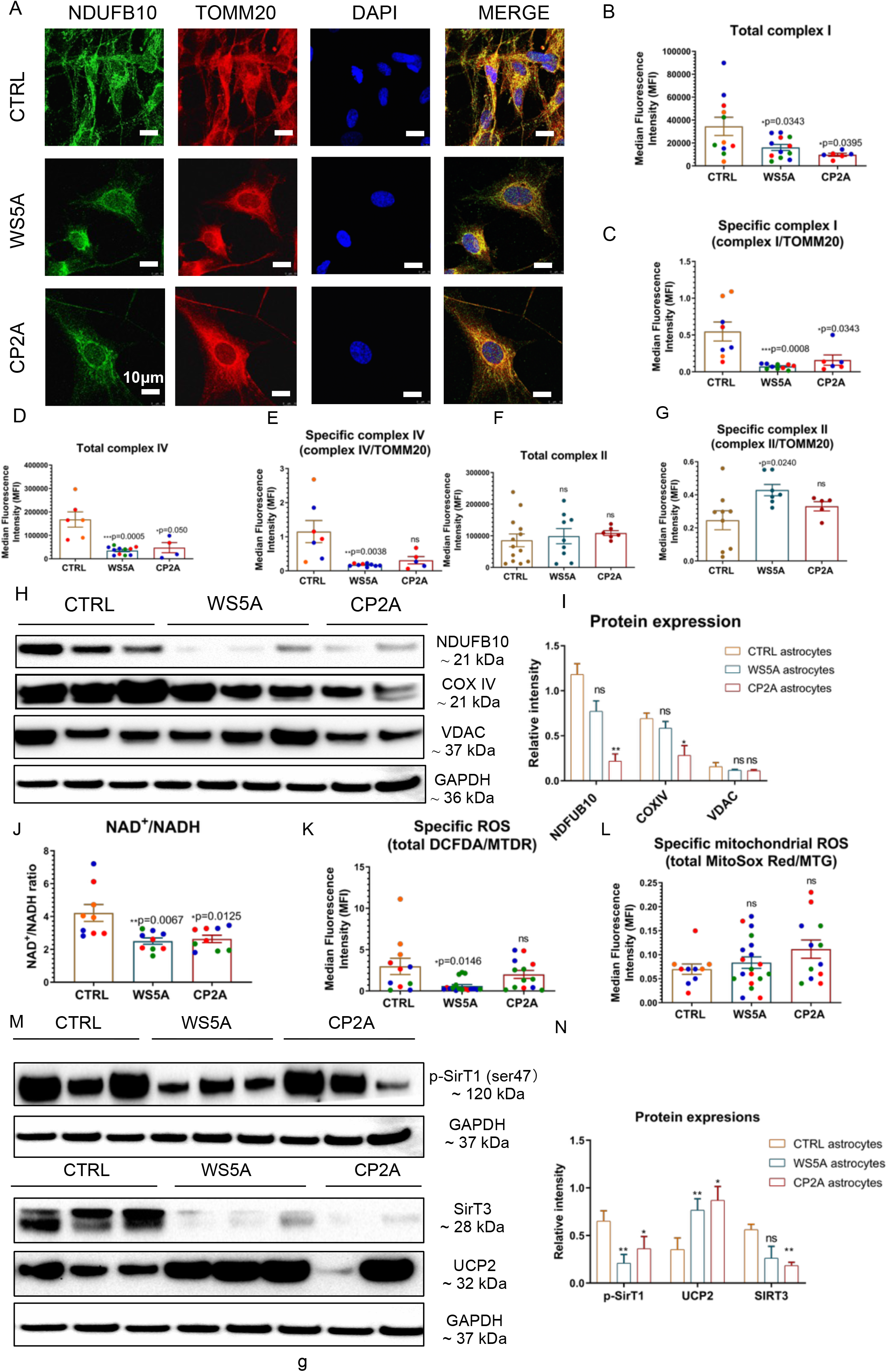
POLG-astrocytes display loss of mitochondrial complex I and IV and disturbed NAD^+^/NADH metabolism. a. Representative confocal images of immunostaining for mitochondrial complex I subunit NDUFB10 (green). Nuclei are stained with DAPI (blue). Scale bar is 10 μm. b-g: Flow cytometric measurements of mitochondrial complex I (b & c), IV (d & e) and II (f & g) protein expression either total level (b, d, f) or specific level (c, e, g) estimated by ratio the total level/TOMM20. h & i. Representative images and quantitation of western blotting for complex I subunit NDUFB10, COX IV, VDAC and GAPDH. j, LC-MS-based metabolomics for quantitative measurement of NAD^+^/NADH ratio. k, Flow cytometric measurements of specific intracellular ROS calculated by total ROS/MTDR using double staining of DCFDA and MTDR. l. Flow cytometric measurements of mitochondrial ROS (mito-ROS) production of the specific level calculated by total mito-ROS/MTG using double staining of MitoSox Red and MTG. m & n. Representative images (m) and quantitation of western blotting for Phospho-SirT1 (Ser47), SirT3, UCP2 and GAPDH (n).

### POLG-astrocytes develop complex I defect that disturbs NAD+/NADH, but not ROS metabolism

Since complex I function is essential for the reoxidation of NADH, and thus for maintaining the NAD^+^/NADH ratio and intermediary metabolism [24], we measured NAD^+^ and NADH levels using LC-MS. As expected, we found a significantly lower NAD^+^/NADH ratio in POLG-astrocytes (Fig. 4j). When comparing to the control, we detected a different change of the absolute levels of NAD^+^ with higher in WS5A lines and lower in CP2A astrocytes (Fig. S19a). We found a significant lower level of total NADH level in CP2A lines comparing to controls, while WS5A astrocyte appeared to be similar with controls (Fig. S19b).

The mitochondrial respiratory chain, particularly complex I, is also considered a major source of reactive oxygen species (ROS) [25,26]. To access ROS production, we used the cell-permeable probe 2’,7’-Dichlorodihydrofluorescein diacetate (DCFDA) and found a deceased ROS level in patient astrocyte versus control astrocytes (Fig. S20a). To measure the specific ROS level per mitochondrial mass, we calculated the mass via MitoTracker Deep Red (MTDR), then we divided total ROS by mitochondrial mass to give specific ROS and, again, found lower level in both patients’ astrocytes with the difference in WS5A lines reached the significance (Fig. 4k). We then investigated the level of mitochondrial ROS using MitoSox and quantified both total and specific ROS as before. We found that the level of total (Fig. S20b) and specific mitochondrial ROS (Fig. 4l) were similar in both patients and controls.

In light of our findings of altered NAD^+^/NADH, we investigated the potential effect on SIRT1, SIRT3 and UCP2 in POLG-astrocytes. We first examined the expression of UCP2, SIRT3 and phosphorylated SIRT1 (Ser47) using western blot analysis and found upregulation of UCP2 expression and reduced SIRT3 and phosphorylated SIRT1 expression in the patient derived astrocytes compared to the control group (Fig. 4m and n). These data suggest that the *POLG* mutations in astrocytes leads to impaired NAD^+^/NADH through down regulation of complex I, which is associated with decreased phosphorylation of SIRT1 and increased UCP2 expression that can potentially explain the loss of MMP.

### Tissue study and cell-specific study display a transcriptomic profiling of A1 specific reactive astrocytic characteristics in POLG-astrocytes

Astrogliosis involves the proliferation and migration of astrocytes following neuronal injury [4,5,27,28]. This process is induced by numerous pathological conditions [29] and can contribute to glial scar formation. Previous work [2] showed that astrogliosis commonly accompanies neuronal loss in POLG-disease, both in acute and chronic lesions. To validate this in the context of our current work, we assessed astrocytic and microglial proliferation in the brain of four patients with POLG-disease. Acute cortical lesions (Fig. 5a (b)) showed severe neuronal loss accompanied by pronounced astrogliosis with GFAP accumulation (Fig. 5a(d)) and microglial activation with increased accumulation of HLA-DR/DP/DQ (Fig. 6a(f)). Unaffected cortex showed no visually appreciable changes in the number of astrocytes and/or microglia (Fig. 5a (a), (c) and (e)) while in chronic affected areas of the brain and spinal cord, severe astrogliosis and microgliosis were present (not shown).

**Figure 5.**
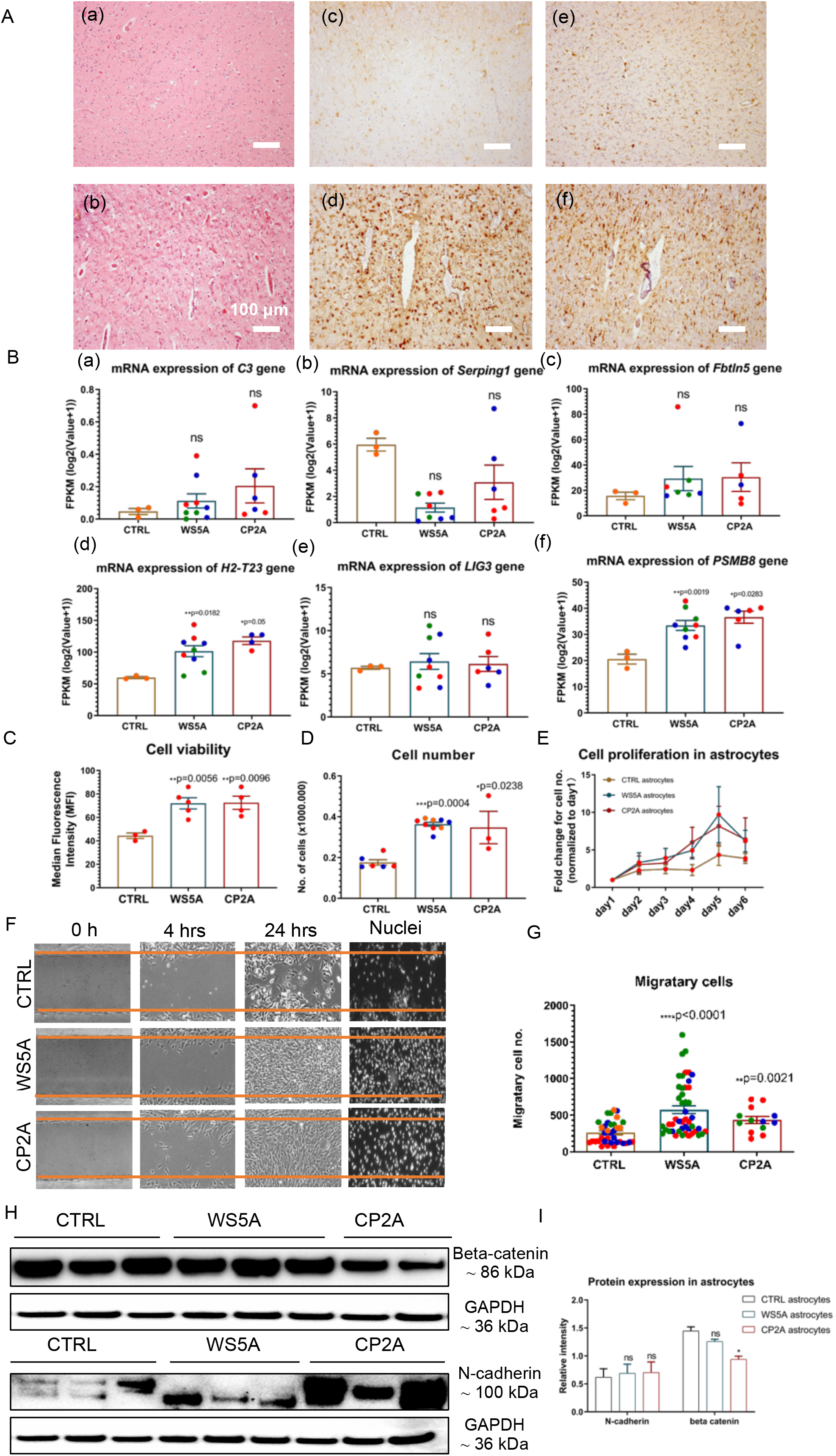
POLG-astrocytes display astrogliosis in brain tissue, a transcriptomic profile of A1 specific reactive characteristics and increased cell viability and migratory ability. a. Brain histology from a POLG patient with p.W748S homozygous mutation. The top row ((a) (c) and (e)) shows unaffected occipital cortex, whereas the lower row ((b) (d) and (f)) shows a cortical strokelike lesion. H&E staining ((a) & (b)), GFAP immunohistochemistry ((c) & (d)) and HLA DR/DP/DQ immunohistochemistry ((e) & (f)) are shown. b. RNA sequencing analysis of mRNA expressions for A1 astrocyte markers. c. Measurement of cell viability using trypan blue staining. d & e. Cell proliferation assays with quantification of cells numbers on day 5 (d) and growth curve from day 0 - day 6 (e). f & g. Representative phase-contrast images of wound healing assay for measurement of cell migration at 0 h, 4 hrs and 24 hrs (f) and quantification of the migrative cell numbers at 24 hrs (g) using DAPI staining for control, WS5A and CP2A astrocytes. Magnification is 100X. h & i. Representative images of western blotting for beta-catenin, N-cadherin and GAPDH and the quantification of their protein expressions.

**Figure 6.**
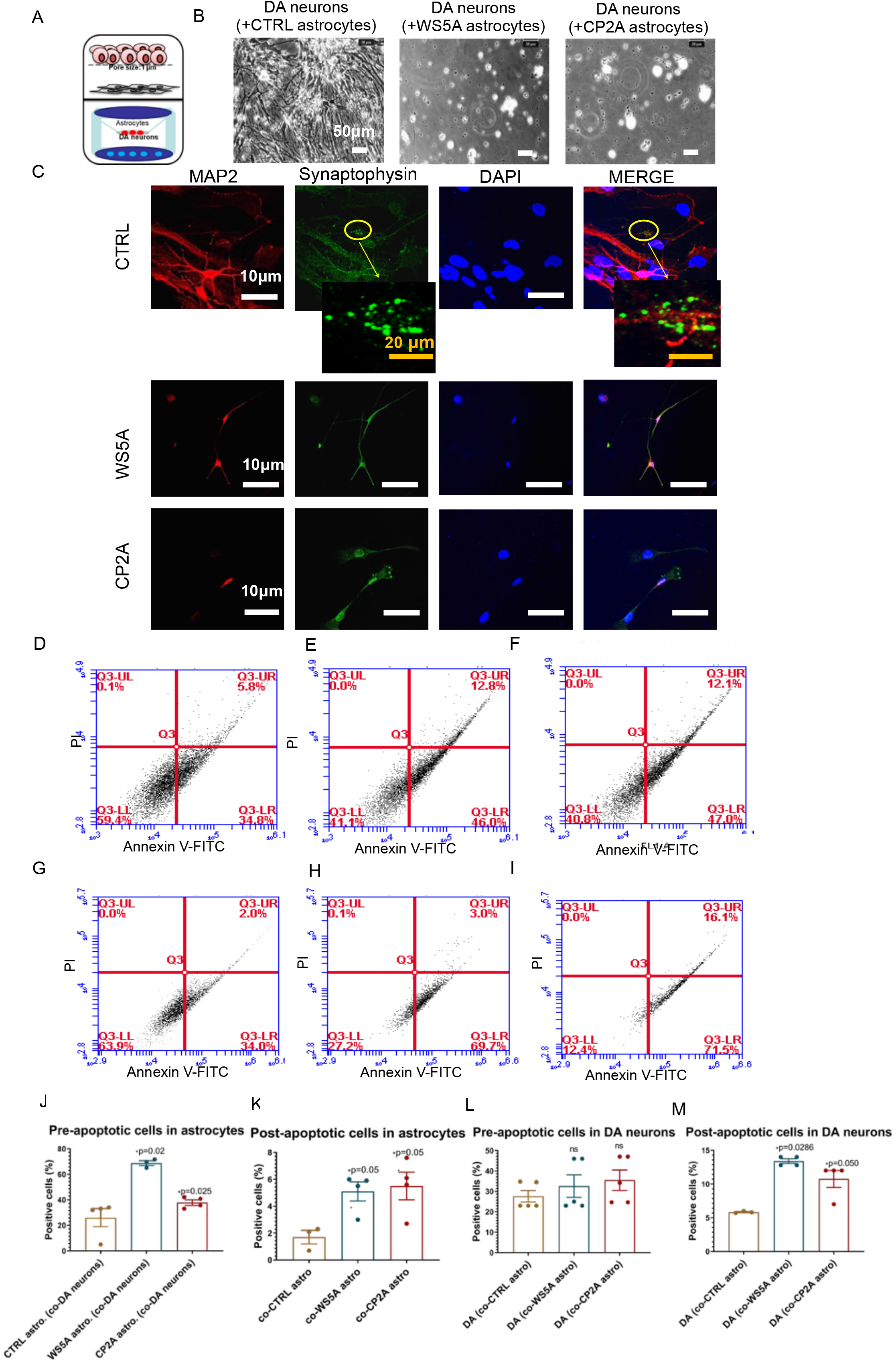
POLG-astrocytes show neurotoxic potential in both direct and indirect neuron/astrocyte co-culture system. a. Illustration of the experimental design for indirect co-culture system with astrocytes and neurons. b. Representative phase-contrast images for DA neurons co-cultured with distinct astrocytes derived from control, WS5A and CP2A iPSCs for 20 days. Scale bar is 50 μm. c. Representative confocal images for double immunostaining of TOMM20 (green) and synaptophysin (red). Second panel showed the images taken from higher magnification from the cells on the first panel. Orange arrow shows the synapse signal. Nuclei are stained with DAPI (blue). White scale bar is 10 μm. Orange scale bar is 20 μm. d - f, Representative flow cytometry plots for the percentage of pro-/early and post-/late apoptotic cells cytometry using the Annexin V-FITC/PI double staining for control (d), WS5A (e) and CP2A (f), astrocytes co-cultured with DA neurons in in-direct co-culture system. g - i. Representative flow cytometry plots for the percentage of pro-/early and post-/late apoptotic cells cytometry using the annexin V-FITC/PI double staining for DA neurons co-cultured with control (g), WS5A (h), and CP2A (i) astrocytes in in-direct co-culture system. j & k. Quantification by calculating the percentage of pro-/early (j) and post-/late (k) apoptotic cells in e-g. l & m. quantification by calculating the percentage of positively stained cells in h-j.

Reactive astrocytes are divided into A1 or A2, where A1 astrocytes are considered neurotoxic while A2 are neuroprotective. A1 astrocytes also lose the ability to promote neuronal survival and outgrowth [5]. We performed RNA-seq analysis on three WS5A clones, two CP2A clones and a panel of control lines including 4 clones from the Detroit 551 and one clone from the AG05836B derived astrocytes. Firstly, we evaluated both A1-, A2- and Oligo2-lineage astrocyte specific mRNA levels and other relevant pathways. Both patient derived astrocytes showed enriched gene expression for A1 astrocytic markers, including *C3*, *Serping1, H2-T23, LIG3* and *PSMB8*, but not *Fbtln5* (Fig. 6b), while not showing any change/perturbation of A2-specific genes such as *CLCF1, TGM1, PTX3, S100A10, EMP1, SLC10A6, TM4SF1* and *B3GNT5* (Fig. S21), nor the Oligo2-lineage astrocyte specific transcripts like *GFAP, HOXB4* and *SLC1A3* (Fig. S22). Several A2-specific transcripts including *SPHK1* (Fig. S21e), *CD109* (Fig. Fig. S21f) and *PTGS2* (Fig. S21g) showed an enrichment, but failed to reach significance. Gene Ontology (GO) enrichment analyses of upregulated DEGs revealed significant enrichment of several disease-related processes including response to stimulus, immune system process, cell proliferation, and cell killing (Fig. S23). In the cell killing pathway, apoptosis was enriched in both WS5A (Fig. S24a) and CP2A astrocytes versus controls (Fig. S25a) with *LMNB2* in WS5A and CP2A (Fig. S24b) and *CTST* in CP2A (Fig. S25b) being the significantly upregulated DEG. KEGG pathway enrichment analysis of down-regulated DEGs revealed enrichment for several signaling pathways (Fig. S26), including the JAK-STAT pathway, which is thought to be active in neuroprotection in reactive astrocytes [30], WNT/beta-catenin pathway and its cross-talk signaling TGF beta pathway, and PI3K-AKT pathway (Fig. S26). Enrichment for pathways that down-regulated axon guidance was also observed in patient derived astrocytes (Fig. S26). These findings demonstrate that A1-like reactive astrocytes are present in POLG-related diseases, raising the possibility that they help to drive neurodegeneration.

### POLG-astrocytes exhibit higher cell proliferation and viability as well as greater cell migration

Having identified a reactive astrocytic gene profile via RNA sequencing analysis, we next assessed a number of phenotypic properties associated with reactive astrocytes. Interestingly, during daily culture maintenance, we observed that POLG-astrocytes showed an increased growth potential compared with control cells. These observations led us to speculate whether the presence of a *POLG* mutation triggered a shift in astrocytes toward a more aggressive, reactive type of cell, which in turn could contribute to neuronal damage. To answer this question, we investigated cell proliferation, viability and migration and analyzed specific markers for reactive astrocytes, including Beta-catenin and N-cadherin. Cell proliferation and viability assays demonstrated significantly increased cell viability (Fig. 5c) and a greater proliferative capacity (Fig. 5d and e) in POLG derived astrocytes compared with controls. We then used a wound healing assay, monitored by optical microscopy, to assess cell migration. Both visually and after quantification, a greater proportion of patient astrocytes cells crossed the area of injury (gap) when compared with controls, suggesting an enhanced migratory capability in POLG-astrocytes (Fig. 5f and g). Western analysis identified higher expression of N-cadherin and a decreased Beta-catenin expression (Fig. 5h and i) in mutant astrocytes than in controls. Overall, these findings suggest that *POLG* mutations are triggering reactive activation of astrocytes.

### POLG-astrocytes exhibit neurotoxic potential in both an indirect co-culture system and in 3D spheroids

In light of our findings suggesting the presence of reactive astrocytes, we asked whether POLG-astrocytes promoted neuronal death. To investigate this, we generated tyrosine hydroxylase (TH) positive dopaminergic (DA) neurons (Fig. S27). POLG-astrocytes and neurons derived from control iPSCs were grown in separate compartments (Fig. 6a), but allowed the exchange of small molecular nutrients. After 20 days of co-culture we assessed the viability of the neurons. We found that DA neurons died when grown with both WS5A and CP2A reactive astrocytes, but not control astrocytes (Fig. 6b). Since astrocytes are involved in synapse formation, we investigated whether POLG-astrocytes also impacted this function. To do this, we used the same control DA neurons and performed double immunostaining for a mature neuron marker, microtubule-associated protein 2 (MAP2) and the synaptic protein synaptophysin (SYP). We observed small synaptic-like vesicles only in the neurons co-cultured with control astrocytes, but not in those with mutant astrocytes (Fig. 6c). This suggests that POLG-astrocytes are either unable to maintain synapses or actively disassemble them, an important area for future investigation.

With the observed decrease in neuronal viability demonstrated above, we next looked at apoptosis. Using a flow cytometry approach, we investigated Annexin V and propidium iodide (PI) levels in astrocytes (Fig. 6d - f) and DA neurons (Fig. 6g - i). We observed increased pre-/early and post-/late apoptotic cell populations in both astrocytes carrying a *POLG* mutation (Fig. 6j and k) and DA neurons co-cultured with POLG-astrocytes (Fig. 6l and m) after 20 days interaction, although the pro-apoptotic cell populations in neurons did not reach significance (Fig. 6l).

We also investigated neuronal toxicity using a three-dimensional (3D) culture system combining neurons, astrocytes, and oligodendrocytes. Briefly, neurons, astrocytes and oligodendrocytes differentiated from control and patient derived NSCs were counted and an equal number plated in hanging drops to generate 3D aggregates/spheroids (Fig. S28). After 20-30 days of co-culture, triple immunofluorescent staining against the neuron marker MAP2, the astrocyte marker GFAP and the oligodendrocyte marker myelin basic protein (MBP) was performed. We observed the three different cell populations only in control spheroids. In patient spheroids, we observed no MAP2 or MBP staining or greatly reduced amounts (Fig. 7a) supporting our observations that *POLG* mutation induces changes in astrocytes that are toxic for both neurons and mature oligodendrocytes. Complement component 3 (C3) is a characteristic and highly upregulated gene in A1 astrocytes [5], therefore, we investigated C3 expression in our astrocytes using immunohistochemistry. We observed GFAP-positive astrocytes that were C3-positive in both patient derived astrocytes (Fig. 7b) and confirmed upregulation of C3 by Western analysis (Fig. 7c and d). Together, these studies suggest that A1 reactive astrocytes are present in POLG-related diseases, where they potentially contribute to the neuronal degeneration and disease pathophysiology

**Figure 7.**
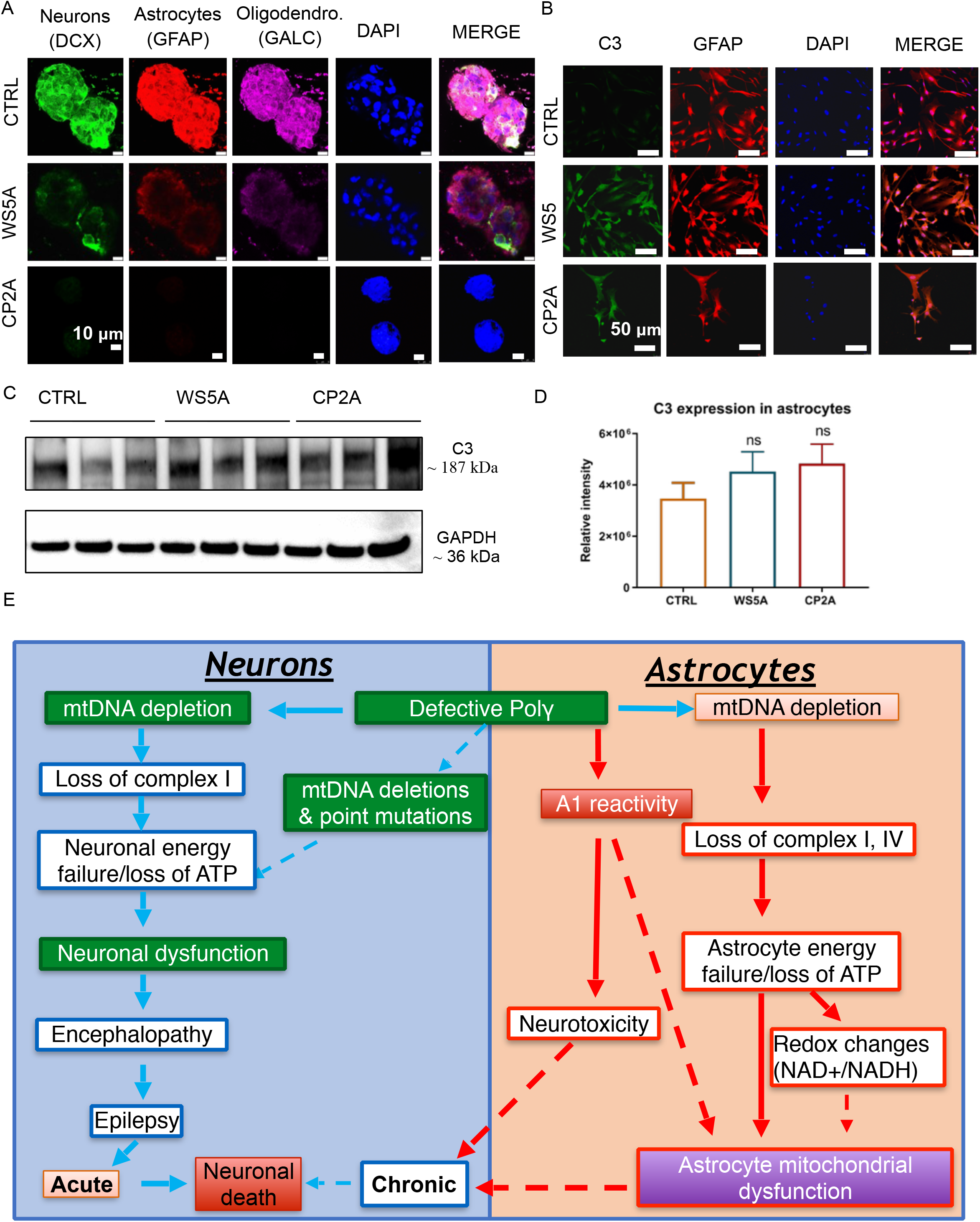
POLG-astrocytes display neurotoxicity in 3D spheroids and a phenotype characteristic of A1-specific reactive astrocytes. a. Representative confocal images of immunostaining for GFAP, DCX and MBP for 3D spheroids composed with astrocytes, oligodendrocytes and neurons from control, WS5A and CP2A iPSCs. Nuclei are stained with DAPI (blue). Scale bar is 5 μm. b. Representative confocal images of immunostaining for C3 for WS5A and CP2A astrocytes. Scale bar is 50 μm. c & d. Representative images (c) and quantitation of western blotting for C3 and GAPDH (d).

## Discussion

### Insert the red ones from version in nature neuronscience

Previous studies, including our own in post-mortem tissue [2] and NSCs [19], have focused on the neuronal consequences of *POLG* mutations. Here, we show, for the first time, that mitochondrial defects including decreased membrane potential, respiratory chain complex loss, lowered ATP levels and mtDNA depletion also occur in iPSC-derived astrocytes. The role of astrocytes in neurodegenerative disease is a rapidly expanding area of interest. For example, mitochondrial defects and oxidative stress were found in cultured SOD1G93A astrocytes and thought to contribute to the motor neuron degeneration in ALS [31]. Further, evidence of astrocytic mitochondrial dysfunction was linked to an early event in Alzheimer disease pathogenesis [12]. More recently, it was shown that mtDNA depletion induced in astrocytes of mice via the inactivation of Twinkle (TwKO mice) produced chronically activated astrocytes and led to an early-onset, spongiotic degeneration of brain parenchyma, microgliosis and secondary neurodegeneration [32]. Our findings show that mtDNA depletion caused by defects in the mtDNA polymerase also generates dysfunctional astrocytes that undergo reactive transition. Our *in vitro* system also allows us to show that these astrocytes are toxic for neurons suggesting that they are actively involved in the disease process.

Astrocytes are the most prevalent cell type in the brain and under normal conditions perform supportive roles for neurons and neuronal circuits. Damage to the central nervous system (CNS) induces the formation of reactive astrocytes that proliferate and generate glial scarring, gliosis, that surround the lesion and separate it from adjacent neural tissue [33–36]. Diseases affecting the CNS can also drive reactive changes [5,37] resulting in either A1 astrocytes, that can kill both neurons and oligodendrocytes, or A2 astrocytes that promote neuronal survival and tissue repair [5,37]. Involvement of A1 astrocytes in various neurodegenerative and inflammatory diseases has been postulated [5]. Here, we show that *POLG* mutations generate A1 like astrocytes with increased cell viability, greater proliferative capacity, and greater ability to migrate. Further, these reactive cells show upregulation of calcium-dependent N-cadherin [38] and a high percentage express C3, a specific reactive astrocyte marker. When we performed RNA sequencing, we found that POLG-astrocytes had significantly up-regulated a cassette of potentially detrimental A1-reactive genes and down-regulated characteristic A2 and Oligo2 genes. This suggests that POLG-astrocytes are already primed to express A1 genes *in vivo*.

Reactive astrogliosis involves both biochemical and structural changes to astrocytes [5] and is driven by various stimuli including hypoxia, inflammation and neurodegeneration. Activated astrocytes show hypertrophy and express a stereotypical array of cytoskeletal proteins, most prominently GFAP. Liddelow et al. [5] reported that induction of a subtype of inflammation-associated reactive astrocytes, that promote death of both neurons and oligodendrocytes, is mediated by activated microglia and others have suggested that this occurs in various neurological diseases [11,17,18]. In the present study, we show that *POLG* mutations induce astrocytic gliosis in vivo, with concurrent microglial activation (Fig. 6a) and in vitro, we show *POLG* mutations are associated with astrocytes having an A1 gene expression profile. Further investigation is necessary to define whether astrocytic reactivity is a primary pathological feature of POLG related disease, and to identify the humeral factors released by astrocytes.

Loss of complex I of the mitochondrial respiratory chain appears to play a crucial role in the neurodegenerative process [39]. Whether this loss is a primary or secondary event is currently, however, unresolved. In Parkinson’s disease, complex I loss was shown to affect the whole brain [40], not just the *substantia nigra* [40,41], suggesting that it may not be the primary cause of neuronal death, but a secondary consequence. We also saw loss of cytochrome c oxidase (COX), complex IV, in POLG astrocytes. This is different to what we found in NSCs, and post-mortem tissue [2]. With regard the latter, COX staining in tissues identifies the largest cells, i.e. in this case the neurons. It is possible, therefore, that any defect in astrocytes would be lost against the background of neuronal staining. Since *POLG* mutations cause mtDNA copy number loss, it is clearly possible that both complexes are lost due to mtDNA depletion. Interestingly, accumulating evidence suggests that complex IV stabilizes the assembly of complex I and that inhibition of complex IV expression can impair complex I assembly in mouse cell lines [42].

In POLG disease, we and others [2,43,44] have demonstrated a complex I defect in tissues and we have confirmed this in iPSC-derived neurons [19]. This loss of complex I is associated with failure to maintain the redox state of the cell, shown by a decreased NAD^+^/NADH ratio, suggesting that in POLG disease, complex I is a primary or at least an important mechanistic event. In addition to the altered redox state, we see impaired mitochondrial energy metabolism shown by lowered ATP production. Emerging evidence implicates NAD^+^ depletion in a wide-range of age-related diseases and neurodegenerative diseases [45–47]. This present study confirms the link between complex I deficiency and altered NAD^+^/NADH ratio and shows that this also occurs in astrocytes. Clearly, these findings also raise the question of whether NAD^+^ supplementation could potentially improve mitochondrial function in POLG related disease.

In conclusion, our current studies demonstrate that *POLG* mutations induce similar mitochondrial changes in astrocytes as are found in POLG neuronal progenitors and post-mortem neurons [19], namely mtDNA depletion, loss of membrane potential and complex I and redox changes. In addition, our current studies provide compelling evidence for the involvement of astrocytes in the disease process initiated by the presence of *POLG* mutations. This allows us to hypothesize that neuronal death in POLG related disease occurs by two mechanisms. Firstly, a process driven by changes in the neuron itself. The loss of ATP with changes in redox potential and ROS production, leads to neuronal dysfunction and chronic neuronal loss. The initiation of seizures in already stressed neurons will exceed the capacity of the neuron to maintain energy metabolism leading to loss of cellular integrity and acute neuronal death. The second, and novel mechanism based on our current work, involves metabolic changes in astrocytes, which induces reactive changes in these cells and leads to chronic neuronal loss (Fig. S29). Finally, we believe our model offers greater tractability than animal models both for investigating mechanisms involved in human disease initiation and progression, and as a system for evaluating therapeutic targets not just for POLG disease, but also in the broader context of neurodegeneration in which complex I deficiency occurs.

## Supporting information

Star Method

## Resource Availability

### Lead Contact

Further information and requests for resources and reagents should be directed to and will be fulfilled by the Lead Contact, Kristina Xiao Liang.

### Materials Availability

This study did not generate new unique reagents.

### Data and Code Availability

The datasets generated and analyzed during the study are included with the Supplemental Information. All other data are available from the corresponding author upon request.

## Method Details

### Astrocytes differentiation

IPSC-derived NSCs were placed on poly-D-lysine (PDL) coated coverslips (Neuvitro, cat.no. GG-12-15-PDL). The following day, the cells were changed into astrocyte differentiation medium, as described in Table S1. The medium was changed every other day for the first week, every two days for the second week and every three days for the third and fourth week. After 28 days of differentiation, the cells were cultured in maturation medium AGMTM Astrocyte Growth Medium BulletKit™ (Lonza, CC-3186) as described in Table S2, for one more month.

### NADH metabolism and ATP measurement using LC-MS analysis

Cells were washed with PBS and extracted by addition of ice-cold 80% methanol followed by incubation at 4°C for 20 min. Thereafter, the samples were stored at −80°C overnight. The following day, samples were thawed on a rotating wheel at 4°C and subsequently centrifuged at 16 000 g at 4°C for 20 min. The supernatant was added to 1 volume of acetonitrile and the samples were stored at −80°C until analysis. The pellet was dried and subsequently reconstituted in a lysis buffer (20 mM Tris-HCl (pH 7.4), 150 mM NaCl, 2% SDS, 1 mM EDTA) to allow for protein determination with BCA assay. The detailed procedure of NADH metabolism and ATP measurement using LC-MS analysis have been described in previous publications [19].

### DNA sequencing for *POLG* mutation

Forward and backward oligonucleotide primers were used to amplify the 7 exons and 13 exons of the *POLG* gene, as reported elsewhere [23,48]. Automated nucleotide sequencing was performed using the Applied Biosystems™ BigDye^®^ Terminator v3.1 Cycle Sequencing Kit (Invitrogen, cat. no.4337454) and analyzed on an ABI3730 Genetic Analyzer with sequencing analyzer software ChromasPro (Technelysium Pty Ltd, Australia). The DNA Chromatogram was aligned with the best matching human sequences in NCBI Trace.

### RNA sequencing

Total RNA was extracted using QIAGEN RNeasy Kit (QIAGEN, 74104). Library preparation was conducted at BGI, Shenzhen, China, following the guide of the standard protocol. Library preparation (BGISEQ-500RS High-throughput sequencing kit, PE50, V3.0, MGI Tech Co, Ltd, Shenzhen, China), hybridization and sequencing were performed according to the manufacturer’s standard procedure provided by BGI (BGI-Shenzhen). The sequencing was performed at BGI-Shenzhen using BGISEQ-500. The sequencing data was filtered using SOAPnuke (v1.5.2) software. The processed FASTQ files were mapped to the human transcriptome and genome using HISAT2 (v2.0.4). The genome version was GRCh38, with annotations from Bowtie2 (v2.2.5). Expression level of the gene was calculated by RSEM (v1.2.12) software.

## Details on the other methods have described in supplementary information

### Ethics approval

The project was approved by the Western Norway Committee for Ethics in Health Research (REK nr. 2012/919); the study was performed in accordance with the Declaration of Helsinki. Tissues were acquired with written informed consent from all patients.

### Competing interests

The authors declare that they have no competing interests.

### Funding

This work was supported by funding from the Norwegian Research Council (project number: 229652), Rakel og Otto Kr.Bruuns legat. G.J.S was partly supported by the Norwegian Research Council through its Centres of Excellence funding scheme (project number: 262613).

### Author’s contributions

K.L and L.A.B contribute to the conceptualization; K.L contribute to the methodology; K.L, A.K, A.C, C.K.K, C.T, Y.H, L.E.H and T.K contribute to the investigation; K.L and A.K contribute to the writing original draft; K.L, C.T, G.J.S and L.A.B contribute to writing review and editing; J.F contributes to the statistical analysis; L.A.B and G.J.S contribute to the funding acquisition; G.J.S and L.A.B contribute to the resources; K.L, L.A.B, and G.J.S contribute to the supervision.

**All authors agree to the authorship.**

## Acknowledgements

We thank members of the Molecular Imaging Centre, Flow Cytometry Core Facility and Genomics Core Facility for their expertise and assistance in confocal imaging and flow cytometry data recording, generating the DNA sequencing data.

## Notes

### Competing Interest Statement

The authors have declared no competing interest.

