## Supplemental Tables and Figures for "Stem cell derived astrocytes with *POLG* mutations and mitochondrial dysfunction including abnormal NAD+ metabolism is toxic for neurons"

**Table S1：  Summary of the information for the cells used in this study.**

| ***Lines*** | ***Status*** | ***Sex*** | ***Age at Biopsy*** | ***Mutation*** |
|---|---|---|---|---|
| HNA | control | F | Fetal | no |
| ESC1 (HS429) | control | F | Embro / 6 days | no |
| ESC2 (HS360) | control | M | Embro / 6 days | no |
| Detroit 551 | control | F | Fetal | no |
| AG05836B | control | F | 44 yrs | no |
| WS5A | POLG disease | F | 44 yrs | c.2243G>C; p.W748S |
| CP2A | POLG disease | M | 49 yrs | c. 1399G>A; p.A467T and c.2243G>C; p.W748S |

**Table S2. Astrocyte differentiation medium.**

| ***Medium/Supplements*** | ***Product no.*** | ***Concentration*** |
|---|---|---|
| DMEM/F-12 + GlutaMAX | Gibco, cat. no. 10565-018 | 500 mL |
| N2 100x | Gibco, cat. no. 17502-048 | 5 mL (1x) |
| B27 50x | Gibco, cat. no. 17504-044 | 10 mL (1x) |
| fibroblast growth factor-2 | Peprotech, cat. no. 100-18B | 4 µg (8 ng/mL) |
| activin A | Peprotech, cat. no. 120-14E | 5 µg (10 ng/mL) |
| heregulin-1ß | Sigma-Aldrich, cat. no. SRP3055 | 5 µg (10 ng/mL) |
| insulin-like growth factor-1 | Sigma-Aldrich, cat. no. I3769 | 100 µg (200 ng/mL) |
| foetal bovine serum | Sigma-Aldrich, cat. no. 12103C | 5 mL (1%) |

**Table S3. Components of the astrocyte maturation medium.**

| ***Medium/Supplements*** | ***Product no.*** | ***Concentration*** |
|---|---|---|
| Astrocyte Basal Medium | Lonza, cat. no. CC-3187 | 500 mL |
| gentamicin | Lonza, cat. no. 17-518Z | 25 mg (50 ng/mL) |
| epidermal growth factor | Gibco, cat. no. PHG0314 | 10 µg (20 ng/mL) |
| ascorbic acid | Sigma-Aldrich, cat. no. A4034 | 500 µg (1 µg/mL) |
| FBS | Sigma-Aldrich, cat. no. 12103C | 15 mL (3%) |
| L-glutamine | Sigma-Aldrich, cat. no. G7513 | 5 mL (1%) |
| insulin | Roche, cat. no. 11376497001 | 1.25 mL (0.25%) |

### Table S4. List of the markers and antibodies and dyes used in this study.

| *Marker* | *Host* | *Company* | *Product no.* | *Application* |
|---|---|---|---|---|
| ***Primary antibodies*** | | | | |
| SOX2 | Rabbit | Abcam | ab97959 | ICC |
| POU5F1 | Rabbit | Abcam | ab19857 | ICC |
| PAX6 | Rabbit | Abcam | ab5790 | ICC |
| NESTIN | Mouse | Santa Cruz Biotechnology | Sc-23927 | ICC |
| NESTIN-PE | Mouse | R&D Systems | IC1259P | FC |
| GFAP | Chicken | Abcam | ab4674 | ICC |
| S100 beta | Rabbit | Abcam | ab196442 | ICC/FC |
| CD44 | Mouse | BD Biosciences | 555476 | FC |
| EAAT1 | Abcam | Abcam | ab416 | ICC |
| DCX | Rabbit | Thermo Fisher | PA5-17428 | ICC |
| Glutamine Synthetase(GluSyn） | Mouse | Abcam | ab64613 | FC |
| NDUFB10 | Rabbit | Abcam | ab196019 | ICC/FC |
| COX IV | Mouse | AbcaM | ab14744 | FC/WB |
| mtTFA (TFAM) | Mouse | Abcam | ab198308 | FC |
| TOMM20 | Mouse | Abcam | ab56783 | ICC |
| GALC | Rabbit | Abcam | ab2894 | ICC |
| TH | Rabbit | Abcam | ab75875 | ICC |
| Synaptophysin | Rabbit | Abcam | ab32127 | ICC |
| Tju 1 | Mouse | Abcam | ab78078 | ICC |
| MAP2 | Chicken | Abcam | ab5392 | ICC |
| SDHA | Mouse | Abcam | ab168536 | FC |
| SDHA | Mouse | Abcam | ab14715 | FC |
| UCP2 | Rabbit | Abcam | 89326 | WB |
| UCP2 | Rabbit | Proteintech | 11081-1-AP | WB |
| Phospho-SirT1 (Ser47) | Rabbit | Cell Signalling | 2314 | WB |
| SirT3 | Rabbit | Cell Signalling | 5490 | WB |
| beta Catenin | Rabbit | Abcam | ab32572 | WB |
| N Cadherin | Rabbit | Abcam | ab76011 | WB |
| GAPDH | Mouse | Abcam | ab8245 | WB |
| C3 | Rabbit | Abcam | ab97462 | WB, ICC |
| ***Secondary antibodies*** | | | | |
| Alexa Flour® 488 | Rabbit | Thermo Fisher Scientific | A11008 | ICC |
| Alexa Flour®594 | Mouse | Thermo Fisher Scientific | A11005 | ICC |
| Alexa Flour®594 | Chicken | Thermo Fisher Scientific | A11042 | ICC |
| ***Dyes*** | | | | |
| DAPI | | Thermo Fisher Scientific | P36962 | ICC |
| MTG | | Invitrogen | M7514 | FC |
| TMRE | | Abcam | ab113852 | FC |
| FCCP | | Abcam | ab120081 | FC |
| DCFDA | | Abcam | b11385 | FC |
| MTDR | | Invitrogen | M22426 | FC |
| MitoSOX™ | | Invitrogen | M36008 | FC |

ICC: Immunocytochemistry staining; FC: Flow cytometry; WB: Western blot,

**Table S5. List of the top 10 DEGs in KEGG metabolism pathway in WS5A astrocytes versus control astrocytes.**

| ***Gene ID*** | ***Gene Symbol*** | ***Qvalue (CTRL-vs-WS5A)*** |
|---|---|---|
| 64131 | *XYLT1* | 1.24E-07 |
| 5742 | *PTGS1* | 5.31E-05 |
| 11343 | *MGLL* | 6.62E-04 |
| 81849 | *ST6GALNAC5* | 9.31E-04 |
| 4881 | *NPR1* | 0.001204153 |
| 80201 | *HKDC1* | 0.001337 |
| 55790 | *CSGALNACT1* | 0.00140215 |
| 117248 | *GALNT15* | 0.002062158 |
| 218 | *ALDH3A1* | 0.002867196 |

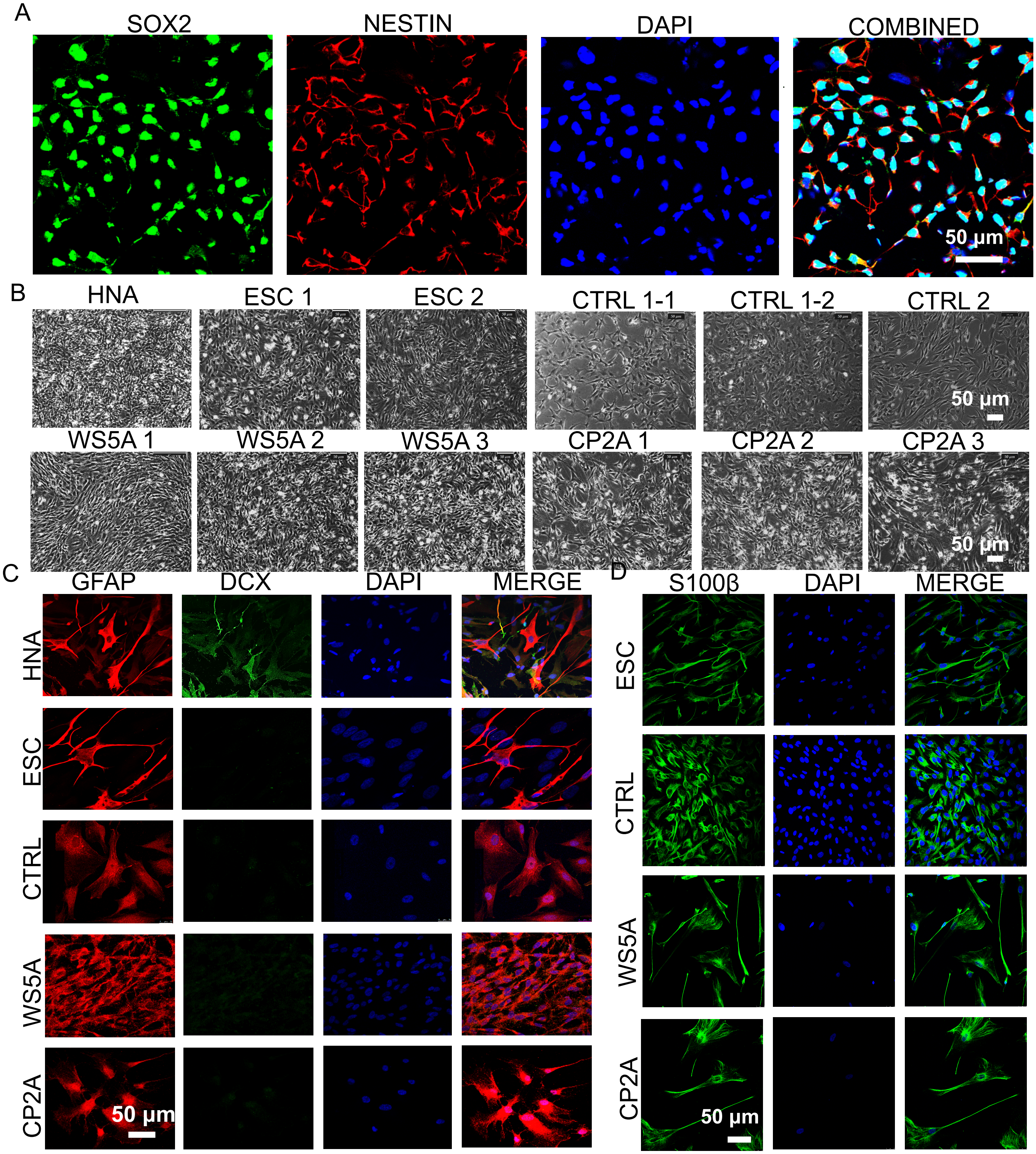


**Fig. S1. Characterization of iPSC-derived NSCs and astrocytes, related to Figure 1 and 2.**
A. Representative confocal images of immunostaining for SOX2 (green) and NESTIN (red) in iPSC-derived NSCs.
B. Representative phase-contrast of NHA, ESC-derived astrocytes and iPSC-derived astrocytes from control and patients carrying homozygous and heterozygous *POLG* mutations (WS5A and CP2A).
C. Representative confocal images of immunostaining for GFAP and DCX in astrocytes.
D. Representative confocal images of immunostaining for S100β in astrocytes. Nuclei are stained with DAPI (blue). Scale bar is 50 µm.

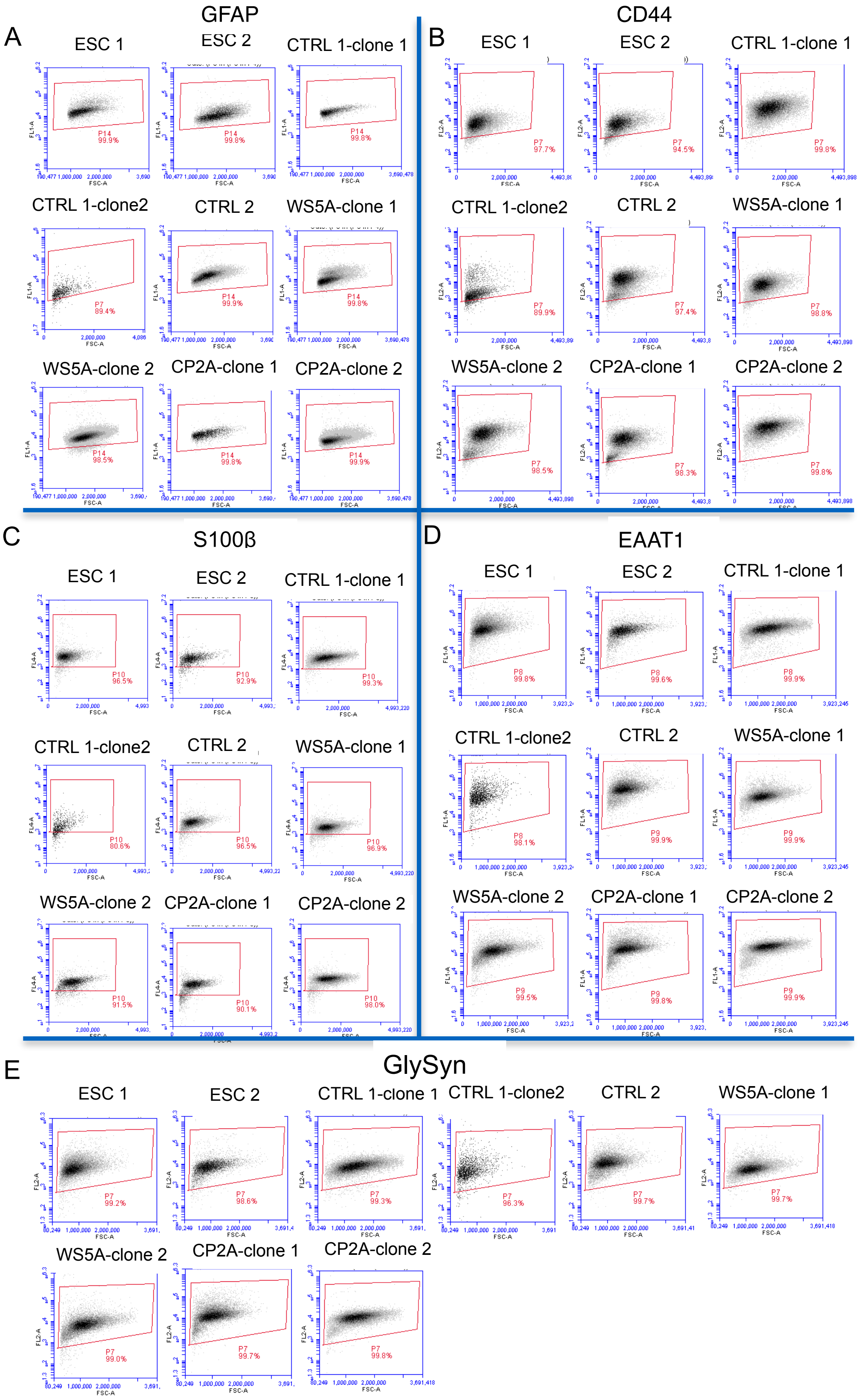


**Fig S2: Representative plots from flow cytometric analysis of the positive cell population stained with astrocyte marker GFAP (A), CD44 (B), S100β (C), EAAT1 (D) and GlySyn (E) in astrocytes, related to Figure 1 and 2.**

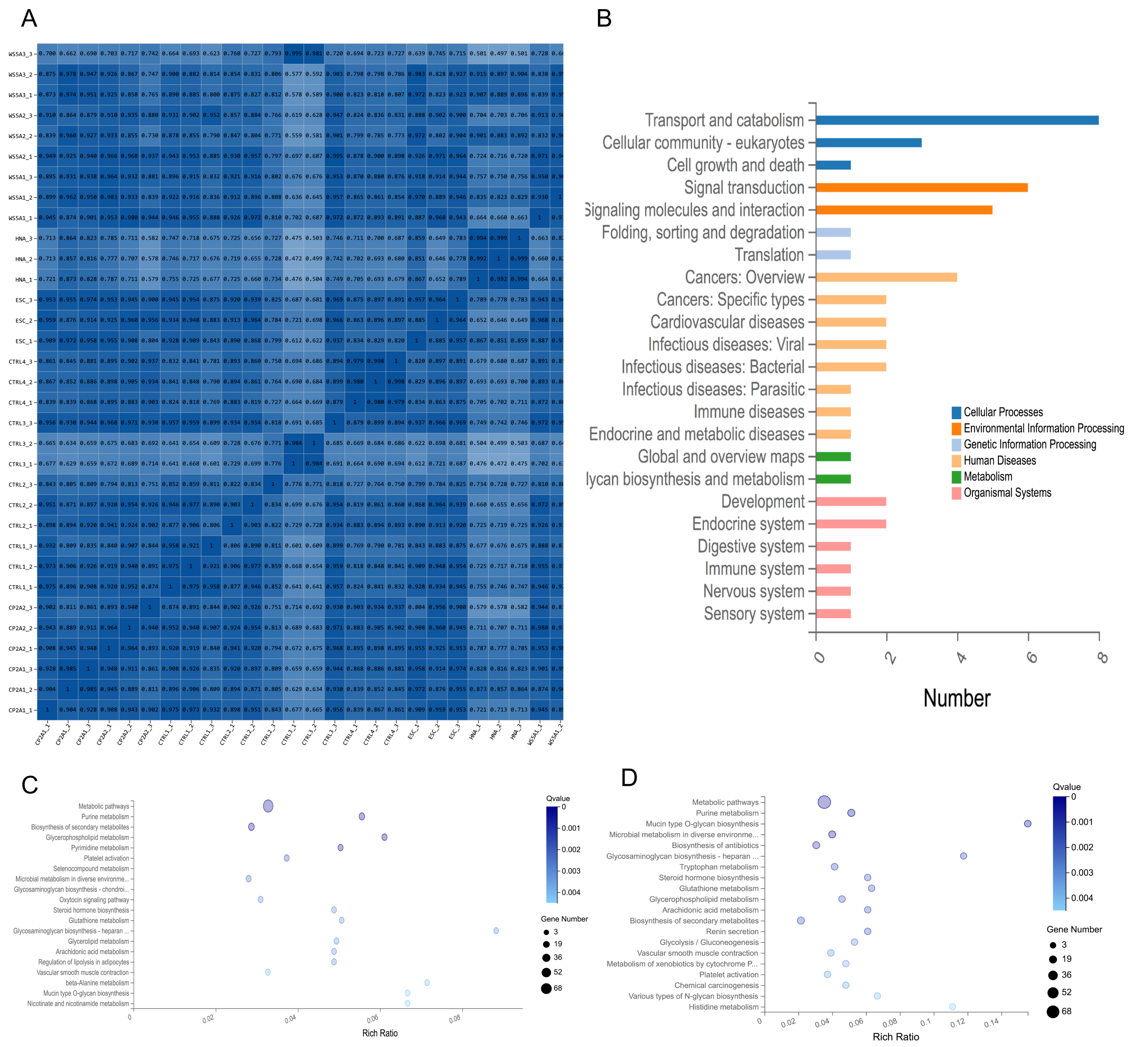


**Fig. S3. RNA sequencing analysis shows a similar transcriptomic profile similar to primary astrocytes and between two patient lines, and *POLG* mutation lead to changes in astrocytes through a metabolic pathway, related to Figure 2.**

A. Correlation heat map of gene expression profiles in individual samples.
B. KEGG pathway classification for DEGs pathway versus control astrocytes.
C. KEGG pathway analysis for up-regulated DEGs in WS5A astrocytes compared to control group.
D. KEGG pathway analysis for up-regulated DEGs in CP2A astrocytes compared to control group.

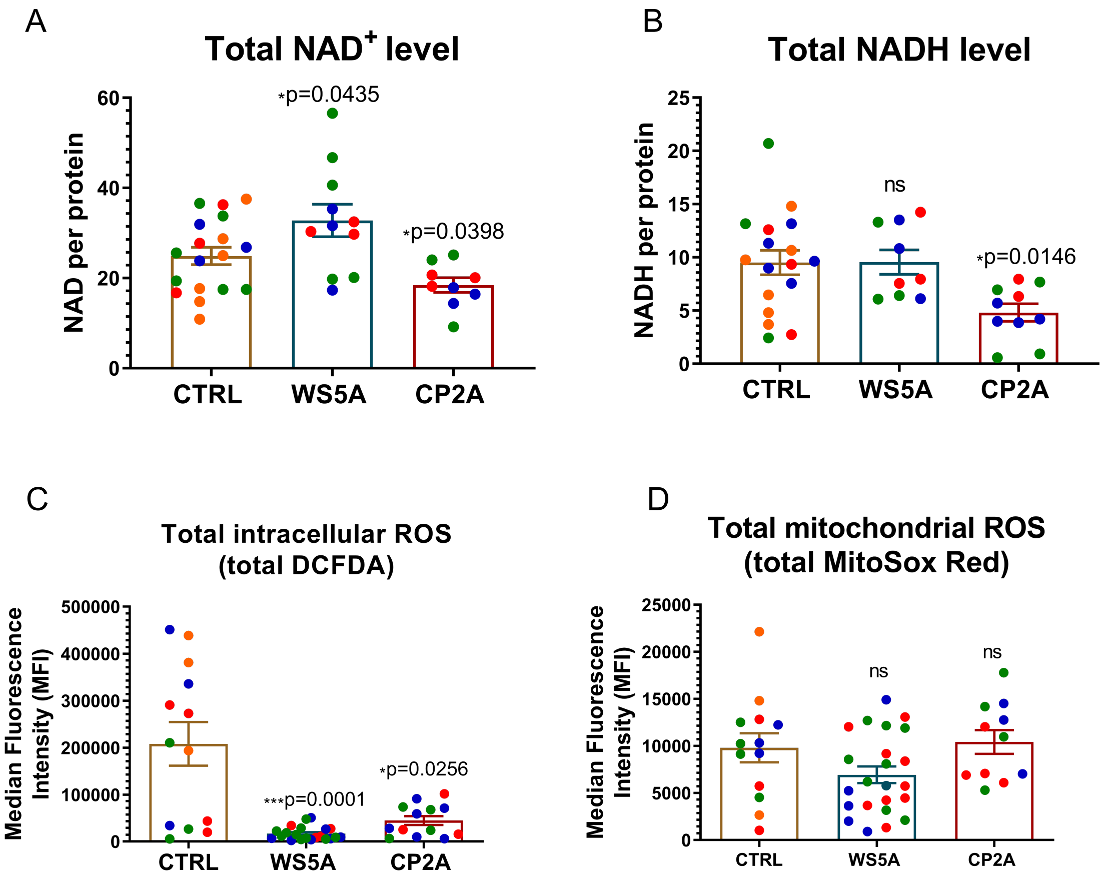


**Fig. S4. Measurement of $NAD^+$ and NADH level using N LC-MS assay and total ROS level using flow cytometry, related to Figure 4.**

A, B. LC-MS-based metabolomics for quantitative measurement of $NAD^+$ (A) and NADH level (B).

C, D. Flow cytometric analysis of intracellular (c) and mitochondrial ROS production (d) at total level in control, WS5A and CP2A astrocytes.

Data information: For the data presented in a - o, red data points represent the data generated from clone #1 from Detroit 551 control, WS5A and CP2A patient lines; blue data points represent the data generated from clone #2 from Detroit 551 control, WS5A and CP2A patient lines; green data points represent the data generated from clone #2 from Detroit 551 control, WS5A and CP2A patient lines and orange data points represent the data generated from AG05836B control. Data are presented as mean ± SEM for the number of samples. Statistical significance was assessed in each patient group compared to the control group. Mann-Whitney U test was used. Significance is denoted for P values of less than 0.05. *P<0.05; ***P<0.001; ns, not significant.

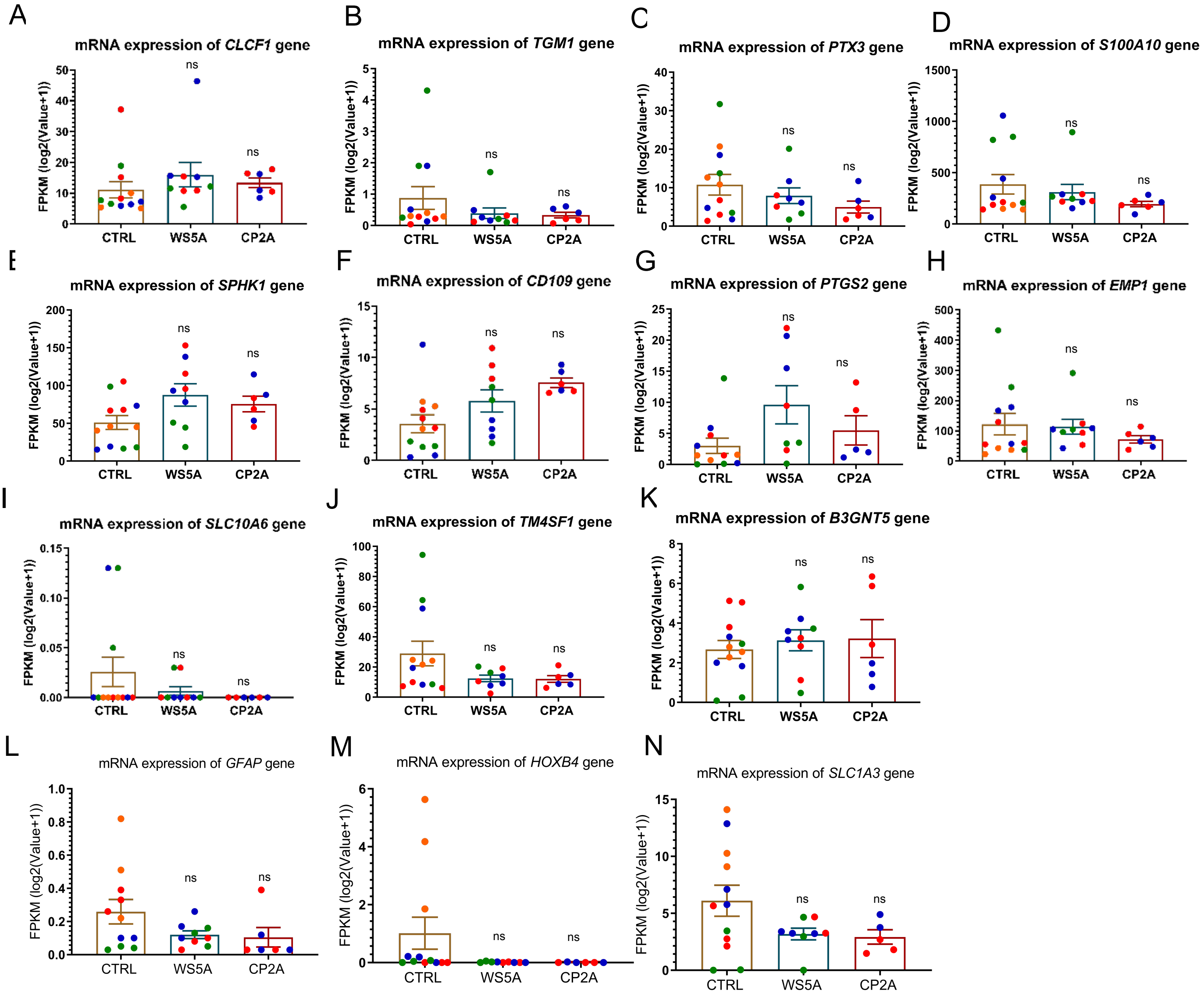


**Fig. S5. RNA-seq. analysis of mRNA expressions for A2 astrocyte (A - K) markers and Oligo2-lineage astrocyte markers (L - N), related to Figure 5.**

Data information: For the data presented in A - N red data points represent the data generated from clone #1 from Detroit 551 control, WS5A and CP2A patient lines; blue data points represent the data generated from clone #2 from Detroit 551 control, WS5A and CP2A patient lines; green data points represent the data generated from clone #2 from Detroit 551 control, WS5A and CP2A patient lines and orange data points represent the data generated from AG05836B control. Data are presented as mean ± SEM for the number of samples. Statistical significance was assessed in each patient group compared to the control group. Mann-Whitney U test was used. Significance is denoted for P values of less than 0.05. ns, not significant.

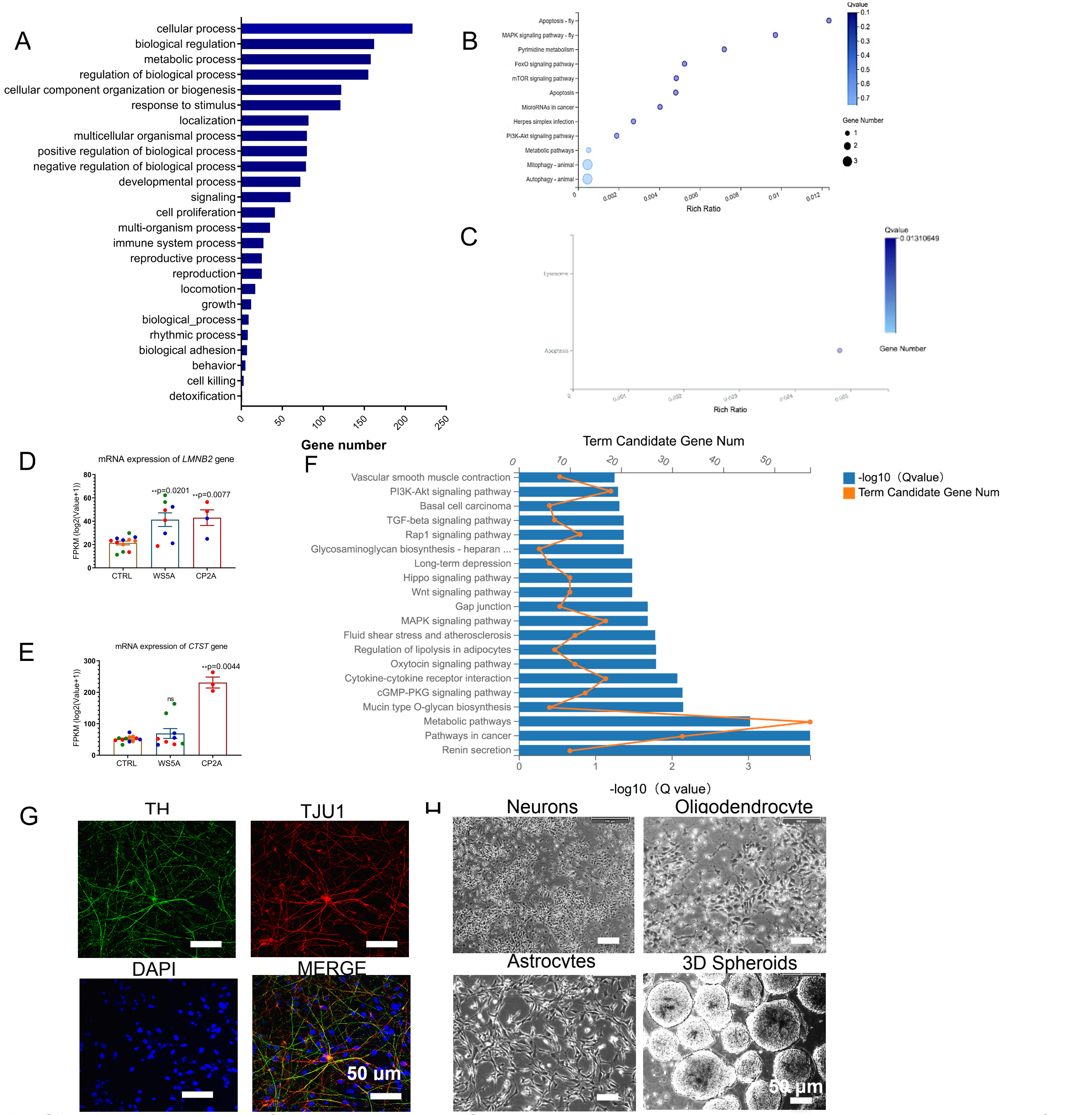


**Fig S6. RNA seq. analysis for up-regulated DEGs in patient astrocytes versus control, characterizaion of DA neurons and images of spheroid formation, related to Figure 6 and 7.**

A.GO-CFP analysis of up-regulated DEGs in WS5A and CP2A astrocytes compared to controls.

B - E. RNA-seq. analysis of KEGG pathway analysis for the DEG enriched in cell killing process in WS5A (B and D) and CP2A (C and E) astrocytes versus controls.

F. Representative confocal images of immunostaining for TH (green) and TJU1 (red) in iPSC-derived DA neurons. Nuclei are stained with DAPI (blue). Scale bar is 50 µm. g. Flow chart of the generation of 3D spheroids using hanging drops with combination of iPSC-derive

G neurons, oligodendrocytes and astrocytes. Scale bar is 50 µm.

Data information: For the data presented in D and E, red data points represent the data generated from clone #1 from Detroit 551 control, WS5A and CP2A patient lines; blue data points represent the data generated from clone #2 from Detroit 551 control, WS5A and CP2A patient lines; green data points represent the data generated from clone #3 from Detroit 551 control, WS5A and CP2A patient lines and orange data points represent the data generated from AG05836B control. Data are presented as mean ± SEM for the number of samples. Statistical significance was assessed in each patient group compared to the control group. Mann-Whitney U test was used. Significance is denoted for P values of less than 0.05. **P<0.01; ns, not significant
