## Supplementary material for "Stem cell derived astrocytes with *POLG* mutations and mitochondrial dysfunction including abnormal NAD+ metabolism is toxic for neurons": Star Method

***STAR★Methods***

***Key Resources Table***

| ***REAGENT or RESOURCE*** | **SOURCE** | **IDENTIFIER** |
| --- | --- | --- |
| ***Anibodies*** | | |
| anti-SOX2 | Abcam | Cat# ab97959, RRID:AB_2341193 |
| anti-POU5F1 | Abcam | Cat# ab19857, RRID:AB_445175 |
| anti-PAX6 | Abcam | Cat# ab5790, RRID:AB_305110 |
| anti-NESTIN | Santa Cruz Biotechnology | Cat# sc-23927, RRID:AB_627994 |
| anti-NESTIN-PE | R&D Systems | Cat# IC1259P, RRID:AB_2151147 |
| anti-GFAP | Abcam | Cat# ab4674, RRID:AB_304558 |
| anti-S100 beta | Abcam | Cat# ab196442, RRID:AB_2722596 |
| anti-CD44 | BD Biosciences | Cat# 555476, RRID:AB_395868 |
| anti-EAAT1 | Abcam | Cat# ab416, RRID:AB_304334 |
| anti-DCX | Thermo Fisher Scientific | Cat# PA5-17428, RRID:AB_10977233 |
| anti-Glutamine Synthetase (GluSyn） | Abcam | Cat# ab64613, RRID:AB_1140869 |
| anti-NDUFB10 | Abcam | Cat# ab196019 |
| anti-COX IV | AbcaM | Cat# ab14744, RRID:AB_301443 |
| anti-mtTFA (TFAM) | Abcam | Cat# ab198308 |
| anti-TOMM20 | Abcam | Cat# ab56783, RRID:AB_945896 |
| anti-GALC | Abcam | Cat# ab2894, RRID:AB_449091 |
| anti-TH | Abcam | Cat# ab75875, RRID:AB_1310786 |
| Synaptophysin | Abcam | Cat# ab32127, RRID:AB_2286949 |
| anti-Tju 1 | Abcam | Cat# ab78078, RRID:AB_2256751 |
| anti-MAP2 | Abcam | Cat# ab5392, RRID:AB_2138153 |
| anti-SDHA | Abcam | Cat# ab168536, RRID:AB_2857979 |
| anti-SDHA | Abcam | Cat# ab14715, RRID:AB_301433 |
| anti-UCP2 | Cell Signalling Technology | Cat# 89326, RRID:AB_2721818 |
| anti-UCP2 | Proteintech | Cat# 11081-1-AP, RRID:AB_2213793 |
| anti-Phospho-SirT1 (Ser47) | Cell Signalling Technology | Cat# 2314, RRID:AB_561516 |
| anti-SirT3 | Cell Signalling Technology | Cat# 5490, RRID:AB_10828246 |
| anti-beta Catenin | Abcam | Cat# ab32572, RRID:AB_725966 |
| anti-N Cadherin | Abcam | Cat# ab76011, RRID:A_1310479 |
| anti-GAPDH | Abcam | Cat# ab8245, RRID:AB_2107448 |
| anti-C3 | Abcam | Cat# ab97462, RRID:AB_10679468 |
| anti-Alexa Flour® 488 | Molecular Probes | Cat# A-11008, RRID:AB_143165 |
| anti-Alexa Flour®594 | Molecular Probes | Cat# A-11005, RRID:AB_141372 |
| anti-Alexa Flour®594 | Molecular Probes | Cat# A-11042, RRID:AB_2534099 |
| ***Chemicals, Peptides, and Recombinant Proteins*** | | |
| DAPI | Thermo Fisher Scientific | P36962 |
| MTG | Invitrogen | M7514 |
| TMRE | Abcam | ab113852 |
| FCCP | Abcam | ab120081 |
| DCFDA | Abcam | b11385 |
| MTDR | Invitrogen | M22426 |
| MitoSOX™ | Invitrogen | M36008 |
| ***Critical Commercial Assays*** | | |
| Lactate Colorimetric/Fluorometric Assay Kit | Abcam | Cat# ab65331 |
| ***Software and Algorithms*** | | |
| [SPSS Statistics 25](https://www.ibm.com/support/pages/downloading-ibm-spss-statistics-25) | IBM | https://www.ibm.com/ |
| GraphPad Prism version 8 for Windows | GraphPad Software, Inc | https://www.graphpad.com/ |
| C6 Plus Workstation Computer and Software | BD Biosciences | https://www.bdbiosciences.com/us/instruments/research/cell-analyzers/bd-accuri/bd-accuri-c6-plus-options/c6-plus-workstation-computer-and-software/p/661391 |
| Image J software | NIH | https://imagej.nih.gov/ij/index.html |
| ChromasPro DNA sequence software | Technelysium Pty Ltd | http://technelysium.com.au/wp/chromaspro/ |
| Image Lab Software | BioRad | https://www.bio-rad.com/en-us/product/image-lab-software?ID=KRE6P5E8Z |
