## Supplemental Tables and Figures for "Stem cell derived astrocytes with *POLG* mutations and mitochondrial dysfunction including abnormal NAD+ metabolism is toxic for neurons"

***Supplementary information***

***Method details***

**Derivation of iPSCs and NSCs generation**

The Norwegian Research Ethics Committee (2012/919) granted ethical approval for the project. Human iPSCs were reprogrammed from fibroblasts as described in our previous publication [1]. As shown in Supplementary Table 1, both POLG and a panel of control iPSCs were used in this study, including two POLG patients, one homozygous c.2243G>C, p.W748S/W748S (WS5A) and one compound heterozygous c.1399G>A/c.2243G>C, p.A467T/W748S (CP2A) patient, control iPSC lines from Detroit 551 (ATCC® CCL 110™), AG05836B fibroblasts (RRID: CVCL_2B58) and two human ESCs controls: the hESC line HS429 (female) and HS360 (male).

All iPSC and ESC lines were maintained in E8 medium (Invitrogen, A1517001) on Geltrex (Invitrogen, A1413302) coated 6-well plates (Thermo Scientific, 140675).

NSCs were generated from the POLG and control iPSC lines by lifting cells into a Chemically Defined Medium (CDM) with epidermal growth factor (EGF) and fibroblast growth factor-2 (FGF-2) both at 100 ng/ml, as described previously [1].

**Astrocytes differentiation**

IPSC-derived NSCs were placed on poly-D-lysine (PDL) coated coverslips (Neuvitro, cat.no. GG-12-15-PDL). The following day, the cells were changed into astrocyte differentiation medium, as described in Supplementary Table 1. The medium was changed every other day for the first week, every two days for the second week and every three days for the third and fourth week. After 28 days of differentiation, the cells were cultured in maturation medium AGMTM Astrocyte Growth Medium BulletKit^TM^ (Lonza, CC-3186) as described in Supplementary Table 2, for one more month.

**DA neuron differentiation**

IPSC-derived neurospheres which were generated from 5 days’ neural induction, were maintained in CDM supplemented with 100 ng/ml FGF8b (R&D systems, 423-F8) over a period of 7 days to initiate DA progenitor induction. The following 7 days, the medium was changed to CDM supplemented with 1 µM purmorphamine (PM) (EMD Millipore, 540220-5MG) and 100 ng/ml FGF8b. Termination of the suspension cultures was performed by dissociating the spheres into single cells by incubation with TrypLE™ Express followed by trituration and subsequent plating into monolayers. The DA neurons were matured in DA medium: CDM supplemented with 10 ng/ml BDNF (PeproTech, 450-02) and 10 ng/ml GDNF (PeproTech, 450-10) on Poly-L-Ornithine (Sigma-Aldrich, P4957) and laminin (Sigma-Aldrich, L2020) coated plates.

**Immunocytochemistry and immunofluorescence (ICC/IF)**

Cells were fixed with 4% (v/v) paraformaldehyde (PFA) and blocked using blocking buffer containing 10% (v/v) normal goat serum (Sigma-Aldrich, G9023) with 0.3% (v/v) Triton™ X-100 (Sigma-Aldrich, X100-100ML). The cells were then incubated with primary antibody solution overnight at 4°C and further stained with secondary antibody solution (1:800 in blocking buffer) for 1 h at RT. NSCs were stained with rabbit anti-PAX6, anti-NESTIN (10c2), anti-SOX2 and mitochondrial respiratory chain complex I subunit anti-NDUFB10. Astrocytes and oligodendrocytes were stained with anti-GFAP, anti-S100 beta and anti-CD44 respectively. The antibodies used for DA neuron staining were anti-TH, anti-Beta III Tubulin and anti-MAP2. The secondary antibodies used were Alexa Fluor® goat anti-rabbit 488 Alexa Fluor® goat anti-mouse 594 and Alexa Fluor® goat anti-chicken 594. After incubation with secondary antibodies, the coverslips were mounted onto cover slides using prolong diamond antifade mountant with DAPI (Invitrogen, P36962).

For staining of neurospheres, spheres were spread directly onto cover slides and left at RT until completely dry and then fixed with 4% (v/v) PFA. After two washes with PBS, the spheres were covered in PBS with 20% sucrose, sealed with parafilm, and incubated overnight at 4°C. The spheres were blocked with blocking buffer for 2 hrs at RT and the primary antibodies were added to the samples overnight at 4°C. After washing the samples for 3 hrs in PBS with a few changes of buffer, incubation with secondary antibodies (as described above) was conducted overnight at 4°C in a humid and dark chamber. Coverslips were mounted using Fluoromount G (Southern Biotech, 0100 01) before imaging was performed using the Leica TCS SP5 or SP8 STED confocal microscope (Leica Microsystems, Germany).

**Mitochondrial volume and MMP measurement**

To measure mitochondrial volume and MMP, cells were double stained with 150 nM MitoTracker Green (MTG) and 100 nM TMRE for 45 min at 37°C. Cells treated with 100 µM FCCP (Abcam, ab120081) was used as negative control. Stained cells were washed with PBS, detached with TrypLE™ Express and neutralized with media containing 10% FBS. The cells were immediately analyzed on a FACS BD Accuri™ C6 flow cytometer (BD Biosciences, San Jose, CA, USA). The data analysis was performed using Accuri™ C6 software.

**L-lactate production measurement**

L-lactate generation was analyzed by colorimetric L-lactate assay kit (Abcam, ab65331) according to the manufacturer’s instructions. End point lactate concentration was determined in a 96-well plate by measuring the initial velocity (2 min) of the balance between NAD^+^ and NADH by lactate dehydrogenase. Immediately following the extracellular flux assay, the plate was measured at OD 450 nm in a microplate reader (VICTOR™ XLight, PerkinElmer).

**Intercellular and mitochondrial ROS production**

Intracellular ROS production was measured by flow cytometry using dual staining of 30 µM DCFDA (Abcam, b11385) and 150 nM MTDR (Invitrogen, M22426), which enabled us to assess ROS level related to mitochondrial volume. Mitochondrial ROS production was quantified using co-staining of 10 µM MitoSOX™ Red Mitochondrial Superoxide Indicator (Invitrogen, M36008) and 150 nM MTG to evaluate ROS level in relation to mitochondrial volume. Stained cells were detached with TrypLE™ Express and neutralized with media containing 10% FBS. The cells were immediately analyzed on a FACS BD Accuri™ C6 flow cytometer.

**NADH metabolism and ATP measurement using LC-MS analysis**

Cells were washed with PBS and extracted by addition of ice-cold 80% methanol followed by incubation at 4°C for 20 min. Thereafter, the samples were stored at -80°C overnight. The following day, samples were thawed on a rotating wheel at 4°C and subsequently centrifuged at 16 000 g at 4°C for 20 min. The supernatant was added to 1 volume of acetonitrile and the samples were stored at -80°C until analysis. The pellet was dried and subsequently reconstituted in a lysis buffer (20 mM Tris-HCl (pH 7.4), 150 mM NaCl, 2% SDS, 1 mM EDTA) to allow for protein determination with BCA assay.

Separation of the metabolites was achieved with a ZIC-pHILC column (150 x 4.6 mm, 5 μm; Merck) in combination with the Dionex UltiMate 3000 (Thermo Scientific) liquid chromatography system. The column was kept at 30°C. The mobile phase consisted of 10 mM ammonium acetate pH 6.8 (Buffer A) and acetonitrile (Buffer B). The flow rate was kept at 400 µL/min and the gradient was set as follows: 0 min 20% Buffer B, 15 min to 20 min 60% Buffer B, 35 min 20% Buffer B. Ionization was subsequently achieved by heated electrospray ionization facilitated by the HESI-II probe (Thermo Scientific) using the positive ion polarity mode, and a spray voltage of 3.5 kV. The sheath gas flow rate was 48 units with an auxiliary gas flow rate of 11 units, and a sweep gas flow rate of 2 units. The capillary temperature was 256°C and the auxiliary gas heater temperature was 413°C. The stacked-ring ion guide (S-lens) radio frequency (RF) level was at 90 units. Mass spectra were recorded with the QExactive mass spectrometer (Thermo Scientific) and data analysis was performed with the Thermo Xcalibur Qual Browser. Standard curves generated for NAD^+^ and NADH were used as reference for metabolite quantification.

**Flow cytometric measurement of TFAM and mitochondrial complexes levels**

Cells were detached with TrypLE™ Express, pelleted and fixed in 1.6% (v/v) PFA (VWR, 100503 917) at RT for 10 minutes, before permeabilization with ice cold 90% methanol. The cells were blocked using a buffer containing 0.3M glycine, 5% goat serum and 1% bovine serum albumin (BSA) in PBS. For TFAM expression, cells were stained with TFAM and TOMM20 separately. For mitochondrial respiratory chain complexes staining, primary antibody anti-NDUFB10 antibody, anti-COX IV antibody and anti-SDHA antibody [2E3GC12FB2AE2] were added, followed by secondary antibody incubation for the samples stained with anti-NDUFB10 antibody and anti-COX IV. The cells were immediately analyzed on a BD Accuri™ C6 flow cytometer.

**Indirect co-culture system**

The transwell chambers with pore size 0.4 µm (Corning, 3450) were used for epithelial-stromal indirect co-culture experiments. iPSC-derived astrocytes were plated onto the insert membranes, while normal iPSC-derived DA neurons were seeded on the bottom of each plate respectively.

**Cell proliferation and viability assay**

A total number of 100.00 cells were seeded in 96-well plates and cultured for 6 days. The number of cells was counted each day and the cell viability was determined with Trypan blue staining.

**Apoptosis detection by flow cytometry**

Apoptotic cell death was measured by flow cytometry using the Annexin V-FITC/PI double staining kit (Invitrogen, cat no. V13242) according to manufacturer's instructions. The number of viable (Annexin negative/PI negative), early apoptotic (Annexin positive/PI negative), and late apoptotic/necrotic (Annexin and PI positive) cells were determined using Accuri™ C6 software.

**Wound healing assay**

A total number of 5000 cells was seeded into ibidi culture-insert chamber (Ibidi, cat no. 81176). After the cells reached 100% confluence, the insert was removed and cultured for further 24 hrs. Cells were observed and images were captured after 0 h, 4 hrs and 24 hrs.

**Western blotting**

Extraction of protein was done using 1X RIPA lysis and extraction buffer (Sigma-Aldrich, R0278) supplemented with Halt™ Protease and Phosphatase Inhibitor Cocktail (Invitrogen, 78444). Protein concentration was determined using BCA protein assay (Thermo Fisher Scientific, 23227). The cell protein was loaded into NuPAGE™ 4-12% Bis-Tris Protein Gels (Invitrogen, NP0321PK2), and resolved in PVDF membrane (Bio-Rad, 1704157) using the Trans-Blot® Turbo™ Transfer System (Bio-Rad, Denmark). Membranes were blocked with 5% non-fat dry milk or 5% bovine serum albumin [BSA] in TBST for 1 h at RT. Membranes were then incubated overnight at 4°C with anti-UCP2, anti-Phospho-SirT1 (Ser47), anti-SirT3 (D22A3), anti-PINK1, anti-PARKIN, anti-LC3B, anti-P62, anti-BNIP3 and anti-GAPDH antibody conjugated to HRP as a control. After washing in TBST, membranes were incubated with donkey anti-mouse antibody or swine anti-rabbit antibody conjugated to horseradish peroxidase secondary for 1 h at RT. Super signal west Pico chemiluminescent substrate (Thermo Fisher Scientific, 34577) was used as enzyme substrate according to manufacturer's recommendations. The membranes were visualized in SynGene scanner (VWR, USA).

**Tissue studies**

Post-mortem examination was performed on brain tissue from 4 POLG patients, including the prefrontal cortex, occipital cortex, cerebellum and spinal cord. The POLG mutations for these patients are 2 patients (AT-1A/44yrs, female and AT-1B/47yrs, male) with A467T/A467T mutation; 2 patients (WS-1A/41yrs, female and WS-3A/43yrs, female) with W748S/W748S mutation. Informed consent was obtained from all subjects and that the experiments conformed to the principles set out in the WMA Declaration of Helsinki and the Department of Health and Human Services Belmont Report. Hematoxylin-eosin (HE) was performed by standard procedures. Immunohistochemistry was performed using antibodies against GFAP, an astrocytic marker, and HLA-DR/DP/DQ, a microglial marker. Antibodies were obtained from DAKO, Glostrup, Denmark.

**DNA sequencing for *POLG* mutation**

Forward and backward oligonucleotide primers were used to amplify the 7 exons and 13 exons of the *POLG* gene, as reported elsewhere [2,3]. Automated nucleotide sequencing was performed using the Applied Biosystems™ BigDye® Terminator v3.1 Cycle Sequencing Kit (Invitrogen, cat. no.4337454) and analyzed on an ABI3730 Genetic Analyzer with sequencing analyzer software ChromasPro (Technelysium Pty Ltd, Australia). The DNA Chromatogram was aligned with the best matching human sequences in NCBI Trace.

**RNA sequencing**

Total RNA was extracted using QIAGEN RNeasy Kit (QIAGEN, 74104). Library preparation was conducted at BGI, Shenzhen, China, following the guide of the standard protocol. Library preparation (BGISEQ-500RS High-throughput sequencing kit, PE50, V3.0, MGI Tech Co, Ltd, Shenzhen, China), hybridization and sequencing were performed according to the manufacturer's standard procedure provided by BGI (BGI-Shenzhen). The sequencing was performed at BGI-Shenzhen using BGISEQ-500. The sequencing data was filtered using SOAPnuke (v1.5.2) software. The processed FASTQ files were mapped to the human transcriptome and genome using HISAT2 (v2.0.4). The genome version was GRCh38, with annotations from Bowtie2 (v2.2.5). Expression level of the gene was calculated by RSEM (v1.2.12) software.

**Statistical analyses**

In order to minimize the phenotypic diversity caused by intra-clonal heterogeneity which is a common issue for iPSC-related studies, multiple clones from each line were included in the all the analyses. Data was presented as mean ± standard error of the mean (SEM) for the number of samples (n≥3 per clone). Distributions were tested for normality using the Shapiro-Wilk test. Outliers were detected using interquartile range (IQR) and Tukey's Hinges test. Mann-Whitney U test was used to assess statistical significance for variables with non-normal distribution, while two-sided Student's t-test was applied for normal distributed variables. One-way ANOVA test was used to test the significance among three groups. Data was analyzed with SPSS software (SPSS v.25, IBM) and figures were produced by GraphPad Prism software (Prism 7.0, GraphPad Software, Inc.). Significance was denoted for P values of less than 0.05.

For RNA sequencing analysis, the set of differentially expressed genes identified from pairwise comparisons was identified using DEseq2 (v1.4.5) package (BGI, Wuhan, China). The number of reads per kilobase per million reads (RPKM) method was used to calculate the modification levels of unique genes. Significantly differentially expressed genes were defined as ones with at least 0.3 FPKM level of expression in at least one of the conditions and a q-value less than 0.05 by Bonferroni test. KEGG (https://www.kegg.jp/) enrichment analysis of annotated different expressed genes was performed by Phyper (https://en.wikipedia.org/wiki/Hypergeometric_distribution) based on Hypergeometric test.

***Data information***

**Figure 1. Generation and characterization of astrocytes derived from ESCs, control and POLG iPSCs.**

Data information: The data points in f represent astrocytes generated from ESC1 (HS429), one clone from Detroit 551 control, 3 different clones from WS5A patient and one clone from CP2A patient iPSCs. The data points in g and h represent astrocytes generated from two ESC lines, one clone from Detroit 551 control, 3 different clones from WS5A patient and 2 clones from CP2A patient iPSCs. Data are presented as mean ± SEM for the number of samples. One-way ANOVA was used for the data presented. Significance is denoted for P values of less than 0.05. **** P<0.0001; ns, not significant.

**Figure 2. Characterization of functional markers in astrocytes derived from ESCs, control and POLG iPSCs.**

Data information: The data points in d and e represent commercial HNA and iPSC-derived astrocytes from two ESC lines, one clone from Detroit 551 control and one clone from AG05836B, one clone from WS5A patient and one clone from CP2A patient iPSCs. Data in d and e are presented as mean ± SEM for the number of samples. One-way ANOVA was used for the data presented. Significance is denoted for P values of less than 0.05. **** P<0.0001; ns, not significant.

**Figure 3. POLG-astrocytes exhibit impaired mitochondrial function, switched glycolysis and mtDNA alteration.**

Data information: For the data presented in b - j, red data points represent the data generated from clone #1 from Detroit 551 control, WS5A and CP2A patient lines; blue data points represent the data generated from clone #2 from Detroit 551 control, WS5A and CP2A patient lines; green data points represent the data generated from clone #2 from Detroit 551 control, WS5A and CP2A patient lines and orange data points represent the data generated from AG05836 control. Data are presented as mean ± SEM for the number of samples. Statistical significance was assessed in each patient group compared to the control group. Student-t test was used for the data presented in b, e, f and j. Mann-Whitney U test was used for the data presented in c, d, g, h and i. Significance is denoted for P values of less than 0.05. *P<0.05; ** P<0.01; *** P<0.001. **** P<0.0001; ns, not significant.

**Figure 4. POLG-astrocytes display loss of mitochondrial complex I and IV and disturbed NAD^+^/NADH metabolism.**

Data information: For the data presented in b - g and j - l red data points represent the data generated from clone #1 from Detroit 551 control, WS5A and CP2A patient lines; blue data points represent the data generated from clone #2 from Detroit 551 control, WS5A and CP2A patient lines; green data points represent the data generated from clone #2 from Detroit 551 control, WS5A and CP2A patient lines and orange data points represent the data generated from AG05836B control. The data points in I and n represent iPSC-derived astrocytes from 2 clones from Detroit 551 control and one clone from AG05836B, 3 clones from WS5A patient and 2-3 clones from CP2A patient iPSCs. Data are presented as mean ± SEM for the number of samples. Statistical significance was assessed in each patient group compared to the control group. Student-t test was used for the data presented in b - g, j and l. Mann-Whitney U test was used for the data presented in I, k and n. Significance is denoted for P values of less than 0.05. *P<0.05; **P<0.01; ***P<0.001; ns, not significant.

**Figure 5. POLG-astrocytes display astrogliosis in brain tissue, a transcriptomic profile of A1 specific reactive characteristics and increased cell viability and migratory ability.**

Data information: For the data presented in b - e and g, red data points represent the data generated from clone #1 from Detroit 551 control, WS5A and CP2A patient lines; blue data points represent the data generated from clone #2 from Detroit 551 control, WS5A and CP2A patient lines; green data points represent the data generated from clone #2 from Detroit 551 control, WS5A and CP2A patient lines and orange data points represent the data generated from AG05836B control. The data points in i represent iPSC-derived astrocytes from 2 clones from Detroit 551 control and one clone from AG05836B, 3 clones from WS5A patient and 2-3 clones from CP2A patient iPSCs. Data are presented as mean ± SEM for the number of samples. Statistical significance was assessed in each patient group compared to the control group. Student-t test was used for the data presented in c. Mann-Whitney U test was used for the data presented in b, d, e, g and i. Significance is denoted for P values of less than 0.05. *P<0.05; **P<0.01; ***P<0.001; ****P<0.0001; ns, not significant.

**Figure 6. POLG-astrocytes show neurotoxic potential in both direct and indirect neuron/astrocyte co-culture system.**

Data information: For the data presented in j and k, the data points represent the astrocyte generated from clone #1 from Detroit 551 control, WS5A and CP2A patient lines co-cultured with DA neurons clone #1 from Detroit 551 control. For the data presented in l and m, the data points represent the DA neurons clone #1 from Detroit 551 control cocultured with astrocyte astrocytes derived from clone #1 from Detroit 551 control, WS5A and CP2A patient lines respectively; n = 4 biologically independent experiments. Data are presented as mean ± SEM for the number of samples. Mann-Whitney U test was used to assess statistical significance in each patient group compared to the control group, and significance denoted for P values of less than 0.05. *P<0.05; ns, not significant.

**Figure 7. POLG-astrocytes display neurotoxicity in 3D spheroids and a phenotype characteristic of A1- specific reactive astrocytes.**

Data information: The data points in c and d represent iPSC-derived astrocytes from 2 clones from Detroit 551 control and one clone from AG05836B, 3 clones from WS5A patient and 3 clones from CP2A patient iPSCs. n = 3 biologically independent experiments. For the data presented in d, red data points represent the data generated from clone #1 from WS5A and CP2A patient lines; blue data points represent the data generated from clone #2 from WS5A and CP2A patient lines; green data points represent the data generated from clone #2 from WS5A and CP2A patient lines and orange data points represent the data generated from AG05836B control. Data are presented as mean ± SEM for the number of samples. Statistical significance was assessed in each patient group compared to the control group. Mann-Whitney U test was used for the data presented in d and e. Significance is denoted for P values of less than 0.05. *P<0.05; **P<0.01; ns, not significant.
